# RGC-specific reversal of lipid peroxidation drives neuroprotection and vision restoration in optic nerve ischemia by targeting GPX4

**DOI:** 10.64898/2026.08.25.747113

**Authors:** Ming Yang, Jie Pan, Shweta Modgil, Rishita Pujari, ChenJie Pan, Anas Alkhabaz, Xiaobai Ren, Liping Liu, Mohammad Ali Shariati, Tazbir Ahmed, Hugo Wu, Roopa Dalal, Yaping Joyce Liao

## Abstract

Nonarteritic anterior ischemic optic neuropathy (NAION) is the leading cause of acute optic nerve-related vision loss in older adults, yet no disease-modifying therapy exists. Although ischemia is a defining feature of NAION, prior therapeutic efforts targeting vascular insufficiency or nonspecific oxidative stress have failed to prevent irreversible retinal ganglion cell (RGC) degeneration, underscoring an unresolved mechanistic gap between ischemic insult and permanent axonal failure. In this endeavour, we identify lipid peroxidation as an important driver of neurodegeneration in NAION. Analyses of human NAION retina, together with a rigorously validated mouse model, demonstrated a remarkable activation of phospholipid peroxidation within the retina following ischemic injury. RGC-specific overexpression of glutathione peroxidase 4 (GPX4), the only known enzyme capable of directly detoxifying phospholipid hydroperoxides within biological membranes, confers striking protection of RGC survival, axonal integrity, and visual function. We further demonstrate that mitochondrial-targeted GPX4 provides superior protection, suggesting mitochondria as a critical locus of lipid peroxidation–driven vulnerability in NAION. Leveraging real-time multiparametric in vivo imaging to directly interrogate axonal metabolism and function, we demonstrate that RGC-specific GPX4 overexpression robustly restores axonal and retinal mitochondrial abundance, improves ATP bioenergetics, and suppresses superoxide stress following optic nerve ischemia. Mitochondria-targeted GPX4 expression further restores axonal transport and retinofugal projections to central visual targets, thereby stabilizing visual pathway connectivity. Notably, these neuroprotective effects are recapitulated by Ebselen, a clinically tested GPX mimetic, identifying lipid peroxide detoxification as a translatable and imaging-validated therapeutic strategy. Collectively, this work establishes ischemia-induced lipid peroxidation as an essential driver of neurodegeneration in NAION and identifies GPX4 as a key molecular determinant of retinal ganglion cell resilience.

**One Sentence Summary:** Lipid peroxidation as mechanism of vision loss in NAION, and targeting GPX4 leads to neuroprotection and visual restoration

## INTRODUCTION

The human retina and optic nerve exist in an intrinsically pro-oxidant environment due to demanding metabolism (1), continuous light exposure, and lipid-rich neural membranes that render them highly susceptible to lipid peroxidation (2, 3). Their membranes are enriched in long-chain polyunsaturated fatty acids (PUFAs), particularly docosahexaenoic acid, which are essential for membrane function yet highly susceptible to free radical-driven oxidation (4, 5). This vulnerability is further exacerbated under ischemic conditions. Reduced perfusion impairs mitochondrial electron transport, increases reactive oxygen species (ROS) production. As a result, it promotes reperfusion-associated oxidative bursts and accelerates membrane lipid damage. Importantly, lipid peroxidation is not merely a biomarker of oxidative stress but an important driver of ferroptosis—an iron-dependent form of regulated cell death triggered by phospholipid hydroperoxide accumulation (6, 7).

Nonarteritic anterior ischemic optic neuropathy (NAION) is the most common cause of acute optic neuropathy in individuals over 50 years of age, presenting as sudden, painless, and often devastating vision loss (8–10). Unfortunately, the contralateral eye is affected in ∼15% of patients within 5 years or in 21% overall (10, 11), leading to severe disability. NAION is strongly associated with systemic metabolic and vascular risk factors, particularly lipid dysregulation (12, 13). NAION is localized to the anterior optic nerve, which intrinsically has high density of mitochondria. Because the anterior optic nerve has high metabolic activity, it is selectively susceptible to high metabolic stress, mitochondrial dysfunction, and ischemia (14). Loss of retinal ganglion cells (RGCs) and axonal degeneration ultimately underlie permanent vision loss in NAION, a disease for which no effective neuroprotective therapy currently exists (11, 15–17).

The most common risk factors for NAION include hyperlipidemia, hypertension, diabetes mellitus, and cardiometabolic conditions, which all contribute to endothelial dysfunction, impaired autoregulation, and microvascular insufficiency of the short posterior ciliary arteries supplying the optic nerve head (12). The most common structural risk factor for NAION includes a tight compartment of the anterior optic nerve and high risk of metabolic stress, which lead to increased reactive oxygen species and impaired mitochondria entry into the axon (18). In the presence of a structurally crowded optic disc (“disc-at-risk”), this lipid-driven vasculopathic milieu increases susceptibility to progression of focal ischemic injury. Beyond vascular compromise, NAION is also associated with systemic inflammatory changes. Plasma immunoprofiling reveals elevated inflammatory mediators in both acute and chronic phases, including IL-1α and CXCL10 in chronic NAION (19). Taken together, a combination of tight compartment, lipid dysregulation, and inflammation leads to endothelial dysfunction. This reinforces ischemic vulnerability and secondary neurodegeneration in NAION.

Prior therapeutic efforts in NAION have largely focused on non-specific neuroprotection, anti-inflammatory strategies, or vascular modulation, none of which have demonstrated consistent clinical benefit (17, 20–22). This suggests that critical downstream mechanisms linking ischemic insult to irreversible RGC degeneration remain insufficiently targeted. Given the lipid richness of the optic nerve and the oxidative stress caused by ischemic insult, lipid peroxidation-driven vulnerability represents a mechanistically grounded yet underexplored pathogenic mechanism in NAION. Previous studies in glaucoma indicate the high risk of lipid peroxidation in the mammalian optic nerve, and glutathione peroxidase 4 (GPX4) is the only known enzyme capable of directly detoxifying phospholipid hydroperoxides within cellular membranes (23–25). Loss or insufficiency of GPX4 activity permits unchecked lipid peroxidation, leading to cell death, whereas augmentation of GPX4 confers robust neuroprotection across retinal and central nervous system contexts (26).

In this study, we examine whether lipid peroxidation is a mechanistic driver of RGC degeneration in NAION and a potential therapeutic target. By integrating analyses of human NAION retina with a validated experimental model of ischemic optic neuropathy, we characterize the spatial and temporal dynamics of lipid peroxidation following NAION. We further examine whether enhancing lipid peroxide detoxification is sufficient to promote RGC survival, axonal integrity, and visual function. Taking advantage of in vivo imaging and bioenergetic analyses, we investigate how oxidative membrane damage impacts retinal and axonal mitochondrial homeostasis. Together, this work establishes lipid peroxidation-driven neurodegeneration as an essential pathogenic driver in NAION. GPX4-centered strategies can serve as disease-modifying interventions with direct translational potential.

## RESULTS

### Increased lipid peroxidation in human and murine ischemic optic neuropathy

To assess whether biomarker of lipid peroxidation is increased in NAION, we quantified two lipid peroxidation end-products: 4-Hydroxynonenal (4-HNE) (27) and malondialdehyde (MDA) (28), as well as the DNA oxidation marker 8-oxoguanine (8-OXOG). Multimodal clinical imaging revealed profound structural and functional deficits in patients with non-arteritic anterior ischemic optic neuropathy (NAION). Color fundus photography and near-infrared reflectance (NIR) imaging demonstrated optic disc edema in NAION eyes compared with healthy controls (Fig. A). The optic disc flavoprotein fluorescence (FPF) maps showed localized functional impairment at the optic nerve head, consistent with visual field loss detected by standard automated perimetry. Enhanced depth imaging optical coherence tomography (EDI-OCT) further confirmed marked optic nerve head swelling in NAION patients. Quantitative analyses demonstrated significant neurodegeneration and visual dysfunction in NAION. Ganglion cell complex (GCC) thickness and retinal nerve fiber layer (RNFL) thickness were markedly reduced in NAION eyes compared with controls (Fig. B-C; both P < 0.0001). Correspondingly, visual field mean deviation was significantly worsened (Fig. D; P < 0.0001), accompanied by a substantial decline in visual acuity (Fig. E; P < 0.0001). In addition, optic disc FPF values were significantly elevated in NAION patients, indicating altered functional mapping at the optic nerve head (Fig. F; P < 0.0001). Together, these findings demonstrate a strong concordance between structural degeneration and functional impairment in human NAION.

In human NAION retina, 4-hydroxynonenal (4-HNE) levels were elevated (Fig. 1G,H), indicating increased lipid peroxidation and sustained oxidative stress within the ischemic optic nerve head. In the mouse NAION model (Fig. S1A,B, Movie 1-3), retinal 4-HNE (Fig. 1I,J) and 8-OXOG (fig. S1C,D) increased significantly at day 21 post-ischemia (both P < 0.001), with a remarkable white laser injury spot (fig.S1B), a significant decrease of the ganglion cell complex (P < 0.001) (fig.S2A,B), a marked decline of visual acuity (P < 0.001) (fig.S2C), and RGC function (P < 0.05) (fig.S2D). Consistently, MDA was elevated in the retina at both time points at both day 1 (fig.S1E), and day 21 (Fig. 1K) post-NAION. In addition to lipid peroxidation, iron serves as an indispensable cofactor driving ferroptotic cell death. However, to our surprise, ferric iron, ferrous iron, and total iron did not increase on day 1 (fig. S3A), and day 21 post-NAION (fig. S3B), arguing against iron-dependent ferroptosis in NAION. Collectively, these findings indicate activation of lipid peroxidation in NAION. As a major end product of ω-6 PUFA oxidation, 4-HNE reflects persistent membrane lipid damage and is consistent with prior evidence linking optic nerve ischemia to mitochondrial dysfunction and ROS accumulation (Fig. 1C). Together, retinal accumulation of reactive lipid aldehydes and systemic expansion of glycerolipid and phospholipid pools indicate disrupted lipid homeostasis in NAION, potentially linking metabolic imbalance to membrane vulnerability of retinal ganglion cell axons.

**Fig. 1.**
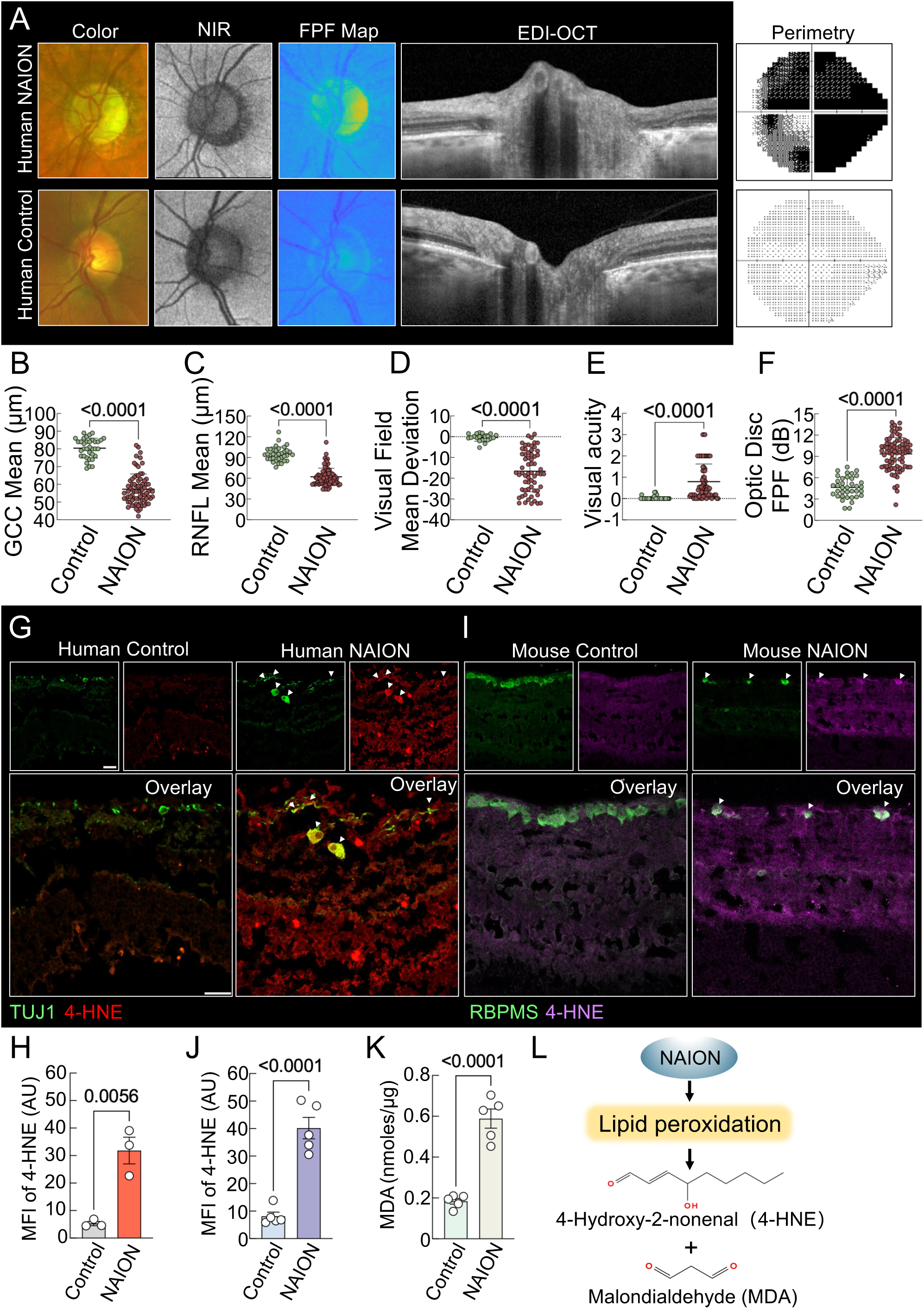
Activated lipid peroxidation in surviving retinal ganglion cells in human and mouse nonarthritic anterior ischemic optic neuropathy (NAION). (A) Representative multimodal retinal imaging from a patient with NAION and a healthy control. From left to right: color fundus photography, near-infrared reflectance (NIR), flavoprotein fluorescence (FPF) map, enhanced depth imaging optical coherence tomography (EDI-OCT), and standard automated perimetry. (B) Quantification of ganglion cell complex (GCC) thickness. NAION eyes exhibit significant thinning compared to controls. (C) Retinal nerve fiber layer (RNFL) thickness is significantly reduced in NAION eyes relative to controls. (D) Visual field mean deviation (MD) demonstrates substantial functional loss in NAION patients. (E) Best-corrected visual acuity is significantly impaired in NAION compared to controls. (F) Optic disc FPF intensity (dB) is significantly increased in NAION, reflecting altered functional mapping at the optic nerve head region. Data are presented as individual data points with mean ± SEM. Statistical significance was determined using a two-tailed unpaired Student’s t-test. (G) Confocal images of human retinal sections showing immunostaining for the neuronal marker Tuj1 together with the lipid peroxidation marker 4-hydroxynonenal (4-HNE). (H) Quantitative analysis of 4-HNE fluorescence intensity in retinas from control and NAION patients. N=3 human retinal sections from 1 NAION patient. Scale bar, 20 µm. Data are presented as mean ± s.e.m.; **p < 0.01, two-tailed t-test. (I) Representative confocal micrographs of mouse retinal sections stained for the retinal ganglion cell marker RBPMS, along with a lipid peroxidation marker 4-HNE. Scale bar, 20 µm. (J) Mean fluorescence intensity of 4-HNE was measured in vehicle-treated and NAION mice 21 days after laser-induced injury. Data was shown as mean ± s.e.m.; **p < 0.01, two-tailed t-test. N=5 control or NAION eyes. (K) Retinal malondialdehyde (MDA) levels were measured in vehicle and NAION mice at days 1 (left panel) and 21 (right panel) following laser injury. Each point represents one sample derived from two pooled retinas. Vehicle (n = 5 mice, 10 retinas) and NAION (n = 5 mice, 10 retinas). (L) A schematic showing that NAION triggers lipid peroxidation, generating MDA and 4-HNE as major end products.

### RGC-specific GPX4 overexpression preserved retinal and optic nerve structure and visual function in animal model of ischemic optic neuropathy

We hypothesize that GPX4 plays a key role in increased lipid peroxidation in NAION because it is the only known enzyme capable of directly detoxifying phospholipid hydroperoxides within cellular membranes (23–25) (Fig. 1C) and overexpression GPX4 is neuroprotective in animal models of optic nerve crush and glaucoma (26). To investigate whether GPX4 overexpression (fig.S4A,B) can dampen lipid peroxidation and rescue RGCs, we performed RGC-specific expression of GPX4—mitochondrial (MitoGPX4) and cytosolic (CytoGPX4) in a mouse model of NAION and performed detailed structural analysis of the retina and optic nerve and comprehensive visual function testing at weekly time points for 3 weeks after NAION. Using optical coherence tomography (OCT), we observed clear and progressive damage in NAION eyes that received a control AAV (Fig. 2B). At day 14 and day 21 after NAION induction, these eyes showed remarkable thinning of the ganglion cell complex (GCC) compared with the untreated contralateral eyes (Fig. 2B,C). The GCC includes the retinal nerve fiber layer (RNFL), ganglion cell layer (GCL), and inner plexiform layer (IPL). This thinning reflects loss of RGCs and their axons, which is typical of NAION. In contrast, eyes treated with GPX4 showed much less GCC thinning, indicating better structural preservation. Among the two forms tested, mitochondrially targeted GPX4 provided slightly stronger protection than cytoplasmically targeted GPX4, as shown in Fig. 2C. Overall, these results show that increasing GPX4 in RGCs reduces structural damage in NAION, with mitochondrial-targeted GPX4 offering a modest additional benefit.

**Fig. 2.**
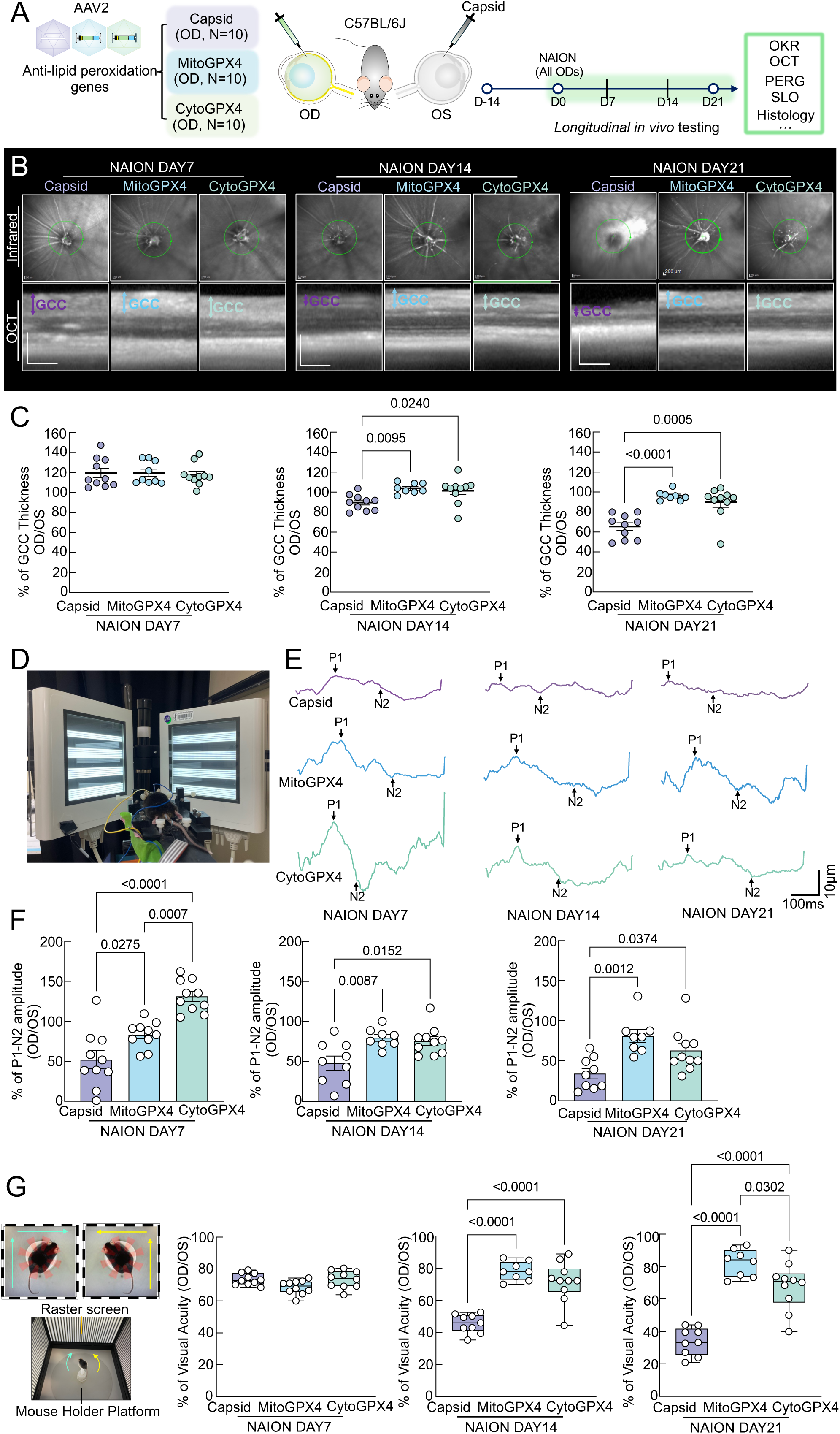
RGC-specific overexpression of GPX4 significantly preserves visual function in the mouse model of NAION. (A) Experimental design illustrating intravitreal delivery of two lipid peroxidation–targeting genes into the right eyes of mice. NAION animals received AAV-Capsid (n = 10), AAV-MitoGPX4, or AAV-CytoGPX4 (n = 10). Retinal structure and function were evaluated longitudinally at 7, 14, and 21 days after NAION induction. The cartoons are optimized with ChatGPT 5.2. (B) Representative in vivo OCT scans of mouse retinas obtained at days 7, 14, and 21 following NAION injury. The ganglion cell complex (GCC), consisting of the retinal nerve fiber layer to inner plexiform layers, is marked by double-headed arrows. Scale bar: 200µm. (C) GCC thickness was assessed by OCT at days 7, 14, and 21 after NAION and expressed as a percentage relative to the corresponding contralateral control eyes. One OCT measurement in the MitoGPX4 group at day 7 was unavailable due to transient cataract formation after anesthesia. AAV-Capsid (n = 10 mice), AAV-MitoGPX4 (n = 9 at day 7, n = 8 at days 14 and 21), and AAV-CytoGPX4 (n = 10 mice). (D) Schematic representation of the pattern electroretinogram (PERG) recording setup used for retinal electrophysiological assessment. The cartoons are optimized with ChatGPT 5.2. (E) Example PERG traces illustrating retinal ganglion cell function. (F) P1–N2 amplitudes of PERG recorded at days 7, 14, and 21 post-NAION were quantified and normalized to contralateral eyes. AAV-Capsid (n = 10 at day 7, n = 9 at days 14 and 21), AAV-MitoGPX4 (n = 10 at day 7, n = 8 at days 14 and 21), and AAV-CytoGPX4 (n = 10 at days 7, 14, and 21). (G) Visual acuity was assessed using the optokinetic response (OKR) and expressed relative to the contralateral eyes. AAV-Capsid (n = 10 at day 7, n = 9 at days 14 and 21), AAV-MitoGPX4 (n = 10 at day 7, n = 8 at days 14 and 21; two mice died at day 14 post-NAION), and AAV-CytoGPX4 (n = 10 at days 7, 14, and 21).

The functional tests were in harmony with structural imaging. Pattern electroretinography (PERG) (Fig. 2D) showed that RGC function was better preserved in eyes treated with GPX4 compared with control-AAV eyes. Again, mitochondrial GPX4 (MitoGPX4) showed slightly stronger protection than cytosolic GPX4 (CytoGPX4). Optokinetic response (OKR) testing, which measures visual acuity, also showed that GPX4-treated eyes maintained better vision. The mitochondrial form provided the greatest functional benefit. Together, these results match the OCT findings. GPX4 overexpression not only preserves retinal structure but also maintains RGC electrical activity (Fig. 2E-F) and visual function in NAION (Fig. 2G, Movie 4).

We also examined retinal tissue after the animals were sacrificed to see whether individual RGCs were preserved. In wholemount retina samples, we counted the number of surviving RGC cell bodies across different regions of the retina. Eyes treated with GPX4 consistently showed many more surviving RGCs than control-AAV–treated eyes. This protective effect was seen in the peripheral, middle, and central areas of the retina (Fig. 3A,B). These in vivo studies suggest that by reducing lipid peroxidation, GPX4 helps protect RGCs throughout the retina in the NAION model. Overall, results from imaging, functional tests, and tissue analysis all point to the same conclusion: increasing GPX4—especially the mitochondrial form—strongly protects RGCs in NAION. It reduces lipid peroxidation–related damage and preserves retinal structure, cell function, and cell survival.

**Fig. 3.**
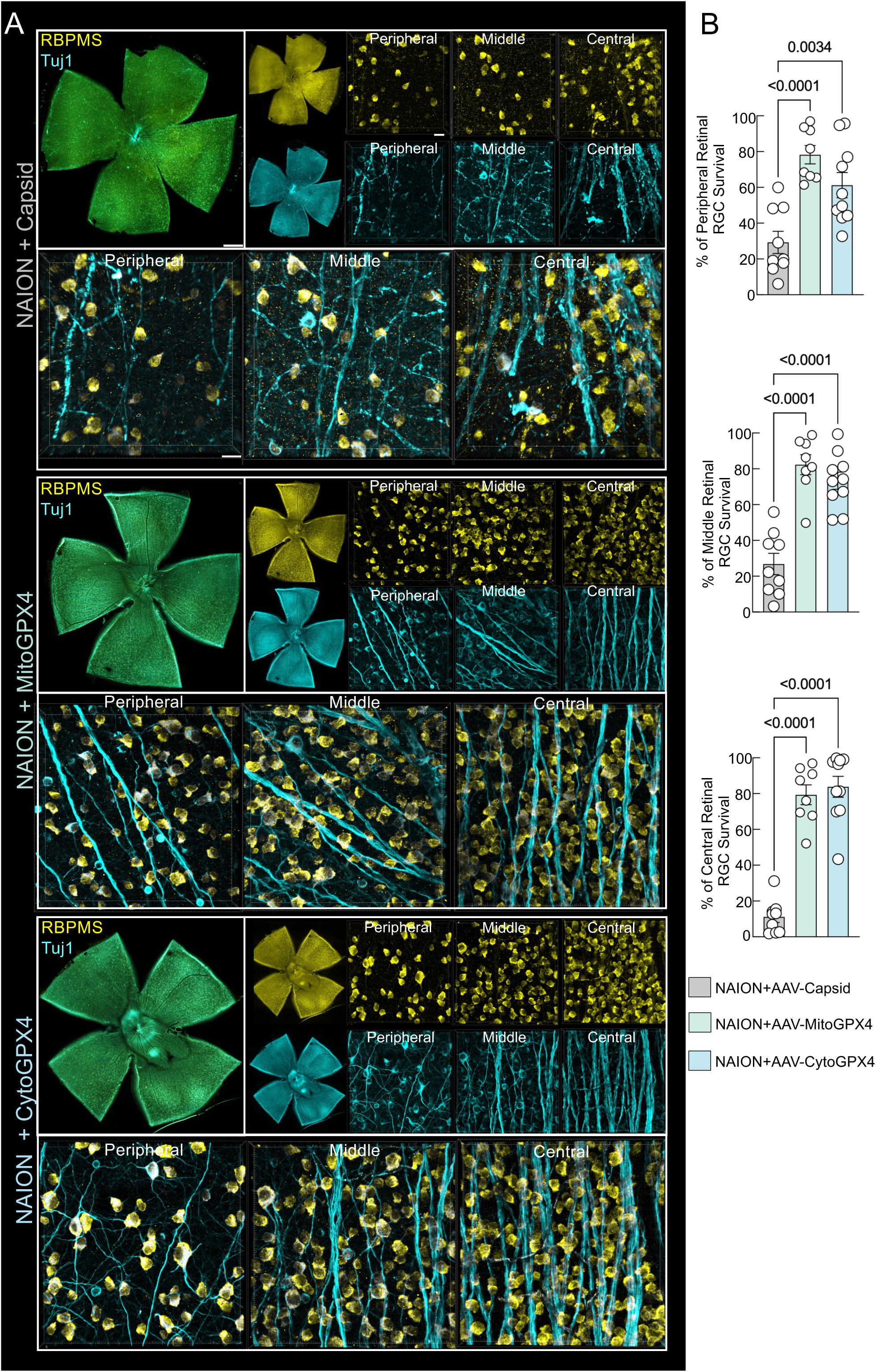
Increased expression of GPX4 confers significant retinal ganglion cell neuroprotection in the NAION model. (A) Whole-mount retinal images illustrating surviving retinal ganglion cells labeled with RBPMS and Tuj1 at 21 days after NAION induction. Low-magnification views of entire flat-mounted retinas are shown (scale bar, 250 µm), together with higher-magnification images from peripheral, mid-peripheral, and central retinal regions (scale bar, 20 µm). Experimental groups include AAV-Capsid (n = 9 mice at day 21 post-NAION), AAV-MitoGPX4 (n = 8 mice at day 21 post-NAION), and AAV-CytoGPX4 (n = 10 mice at day 21 post-NAION). (B) Bar graphs showing quantification of the survival percentages of peripheral, mid, and central RGCs.

### RGC-specific GPX4 overexpression robustly restores axonal and retinal mitochondrial abundance, ATP bioenergetics, and suppresses superoxide stress in vivo

To decipher the mechanism of GPX4-mediated neuroprotection, we take advantage of the confocal scanning laser ophthalmoscope (cSLO)-based in vivo imaging strategy that enables real-time, simultaneous quantification of retinal mitochondrial abundance, ATP bioenergetics, and mitochondrial superoxide stress in the retina (Fig. 4A). Leveraging MitoTracker Green(29), iATPSnFR2(30), and MitoSOX (31), this optimized platform permits longitudinal tracking of mitochondrial dysfunction and metabolic collapse in the living mouse retina, creating a translationally relevant tool for testing neuroprotective and disease-modifying therapies. Our real-time analysis revealed that NAION insult precipitates a catastrophic decrease of mitochondria abundance (Fig. 4B,E), bioenergetic failure, characterized by a profound depletion of intra-retinal RGC axonal ATP (Fig. 4C,F), and a concomitant decrease in mitochondrial superoxide flux (Fig. 4D,G) at day 7 after NAION insult. Remarkably, RGC-specific GPX4 overexpression acted as a metabolic safeguard; it not only restored the mitochondrial abundance, reinvigorated axonal ATP reserves but also successfully quenched mitochondrial oxidative stress, restoring superoxide levels to homeostatic baselines. Remarkably, GPX4 also significantly quenched the activated lipid peroxidation (Fig. 4H). These findings collectively demonstrate that GPX4 does more than prevent lipid peroxidation-it effectively reprograms the mitochondrial landscape, sustaining the energetic demands required for axonal survival under NAION. By enabling real-time, multiparametric imaging of axonal metabolism in vivo, we provide a diagnostic window into early RGC dysfunction, converting molecular neurodegeneration into a quantifiable imaging phenotype for NAION.

**Fig. 4.**
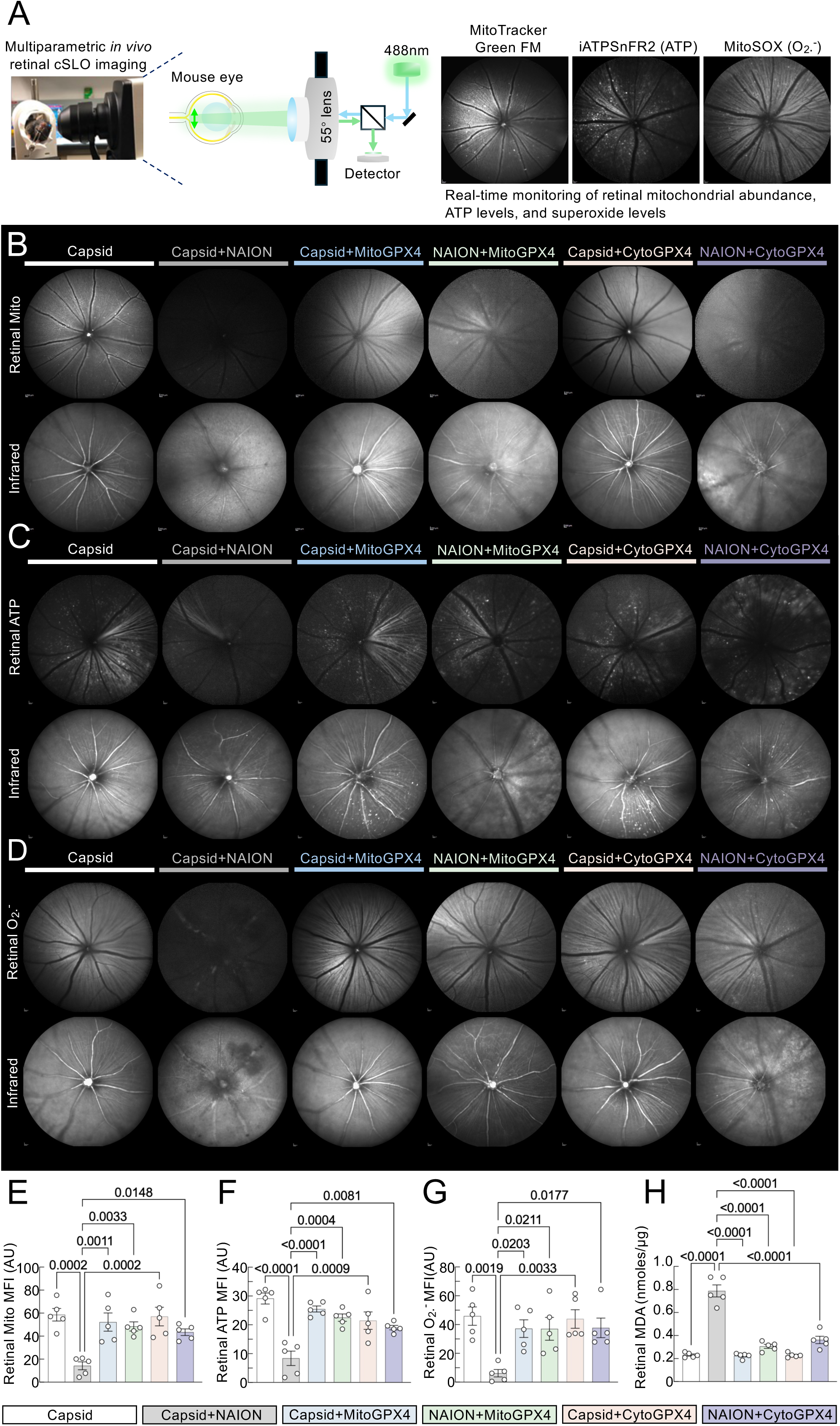
GPX4 restores axonal and retinal mitochondrial abundance, bioenergetics, and suppresses lipid peroxidation in retinal ganglion cells after experimental NAION. (A) Illustration showing a method of multiparametric in vivo imaging to visualize mitochondria abundance, ATP levels, and superoxide levels. (B) Mitochondria abundance in the RGC were monitored by intravitreal injection of Mitotracker in AAV-Capsid (n = 5 mice), AAV-Capsid + NAION (n = 5 mice), AAV-Capsid + AAV-MitoGPX4 (n = 5 mice), NAION + AAV-MitoGPX4 (n=5 mice), AAV-Capsid+AAV-CytoGPX4 (n = 5 mice), and NAION+AAV-CytoGPX4 (n=5 mice) groups. (C) ATP in the RGC was monitored by intravitreal injection of AAV-iATPSnFR2 in AAV-Capsid (n=5 mice), AAV-Capsid + NAION (n = 5 mice), AAV-Capsid + AAV-MitoGPX4 (n = 5 mice), NAION + AAV-MitoGPX4 (n = 5 mice), AAV-Capsid+AAV-CytoGPX4 (n = 5 mice), and NAION+AAV-CytoGPX4 (n = 5 mice) groups. (D) Superoxide in the RGC was monitored by intravitreal injection of MitoSOX in AAV-Capsid (n = 5 mice), AAV-Capsid + NAION (n = 5 mice), AAV-Capsid + AAV-MitoGPX4 (n = 5 mice), NAION + AAV-MitoGPX4 (n = 5 mice), AAV-Capsid+AAV-CytoGPX4 (n = 5 mice), and NAION+AAV-CytoGPX4 (n = 5 mice) groups. (E) Quantification of mean fluorescent intensity (MFI) of mitochondria in the RGC in different groups. (F) Quantification of MFI of ATP in the RGC in different groups. (G) Quantification of MFI of superoxide in different groups. (H) MDA levels of the retina in different treatments were determined. AAV-Capsid (n = 5 mice), AAV-Capsid + NAION (n = 5 mice), AAV-Capsid + AAV-MitoGPX4 (n = 5 mice), NAION + AAV-MitoGPX4 (n = 5 mice), AAV-Capsid+AAV-CytoGPX4 (n = 5 mice), and NAION+AAV-CytoGPX4 (n = 5 mice). Data are presented as means ± s.e.m, one-way ANOVA with Tukey’s multiple comparisons test. Data are presented as means ± s.e.m, one-way ANOVA with Tukey’s multiple comparisons test.

### RGC-specific overexpression of mitochondrial GPX4 restores axonal transport and retinofugal projections after optic nerve ischemia

RGC axons form the essential link transmitting retinal visual signals to central brain regions, and their degeneration in NAION culminates in permanent vision loss due to limited regenerative capacity (32). Therefore, protecting the integrity of RGC axons is critical for vision preservation. In addressing the challenge of protecting RGC axons and their projections under NAION, we employed a targeted genetic approach to overexpress MitoGPX4 specifically in RGCs. Because mitochondrial-targeted GPX isoform overexpression confers more robust neuroprotection than cytosolic GPX, we specifically examined the MitoGPX4 isoform to assess eye-brain connectivity alterations after NAION and to delineate the neuroprotective effects of GPX4 overexpression along the visual pathway. We then used intravitreal cholera toxin subunit B (CTB) tracing to quantify the integrity of the retino-geniculate and retino-collicular pathways, examining brain targets LGN and SC at 21 days post-injury (Fig 5A, Movies 5-7). Baseline comparisons showed that NAION severely disrupts axonal transport, as evidenced by substantial reductions in CTB labelling within central targets. Quantitatively, labelling intensities in SCN, LGN, and SC fell to approximately 53%, 47%, and 53% of uninjured control levels, respectively, indicating broad attenuation of axonal connectivity (Fig 5B–D). When we increased mitochondrial GPX4 in RGCs, the axons connecting the retina to the brain were largely preserved. Using CTB labelling to track these axons, we found that the signal recovered to about 78% of normal levels in the SCN, 71% in the LGN, and 80% in the SC (Fig. 5A–D). This shows strong protection of both the retino-geniculate and retino-collicular visual pathways. These findings suggest that mitochondrial GPX4 in RGCs helps maintain axon structure and axonal transport, which are essential for sending visual information from the eye to the brain. This protective effect goes beyond simply reducing lipid peroxidation. Together, the data support a model in which RGC mitochondrial redox restoration via GPX4 maintains axonal health and preserves central visual circuitry after NAION, thereby potentially sustaining residual vision during disease.

**Fig. 5.**
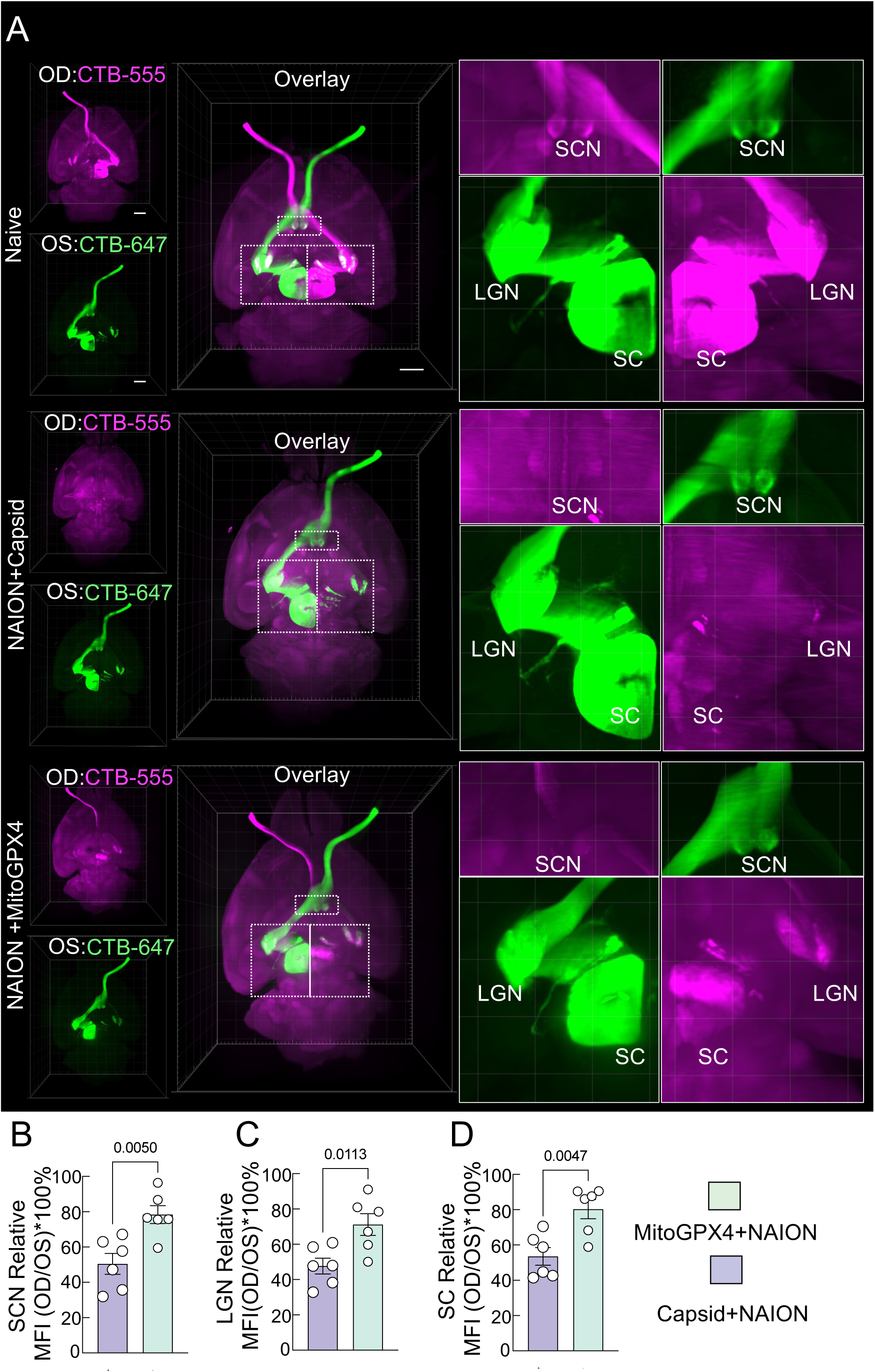
Increased expression of GPX4 provides substantial neuroprotection across the visual pathway in a NAION model. (A) Representative 3D images of the eye and brain were reconstructed following cleared-tissue imaging using a Light-sheet Ultramicroscope II. The visual transportation is labeled by intravitreal injection of cholera toxin B (CTB)-555 and CTB-647 to each eye. (B) Quantification of relative fluorescent intensity of CTB signal in superior colliculus (SC), AAV-Capsid + NAION (n = 6 mice), AAV-CytoGPX4+NAION (n = 6 mice). (C) Quantification of relative fluorescent intensity of CTB signal in the lateral geniculate nucleus (LGN), AAV-Capsid + NAION (n = 6 mice), AAV-CytoGPX4+NAION (n = 6 mice). (D) Quantification of relative fluorescent intensity of CTB signal in the SC, AAV-Capsid + NAION (n = 6 mice), AAV-CytoGPX4+AION (n = 6 mice). Scale bar: 500 µm. Data are presented as means ± s.e.m, two-tailed t-test.

### GPX mimetic Ebselen prevents neurodegeneration and preserves visual functions in a mouse NAION model

We show that RGC-specific AAV2-mediated GPX4 overexpression leads to robust neuroprotection, so the next step is to translate these exciting findings to patients. Given the potential challenges of development of gene therapy as treatment for human disease, we examined the effect of a GPX mimetic small molecule called Ebselen, which has been assessed in clinical trials for ischemic stroke(33), hearing loss, and tinnitus(34). Ebselen is a lipophilic radical-trapping antioxidant (35–38) and a potent small molecule glutathione peroxidase enzyme mimic, which is designed based on the glutathione peroxidase-active sites (39). To investigate the efficacy of Ebselen as treatment for NAION, we first improved the formulation of the drug by improving solubility of this hydrophobic drug (Methods) and then assessed its efficacy as IP injection in mice within 30 min after NAION (Fig. 6A). To assess structural impact of Ebselen treatment, first, we performed serial in vivo retinal imaging using near infrared OCT and segmentation of the retinal thickness, which revealed that Ebselen-treated eyes maintained substantially thicker GCC (RNFL, GCL, IPL) at days 14 and 21 compared with control-treated NAION eyes, which showed pronounced GCC thinning (Fig. 6B-D). Functionally, Ebselen treatment after NAION preserved RGC electrophysiology as measured by PERG (Fig. 6F-H), and maintained visual acuity confirmed by OKR (Fig. 6I-K, Movie 8). Postmortem analyses confirmed higher survival of RGC somas (Fig. 7A,B) in the retina across peripheral to central regions and greater preservation of RGC axons in the optic nerves (Fig. 7C,D).

**Fig. 6.**
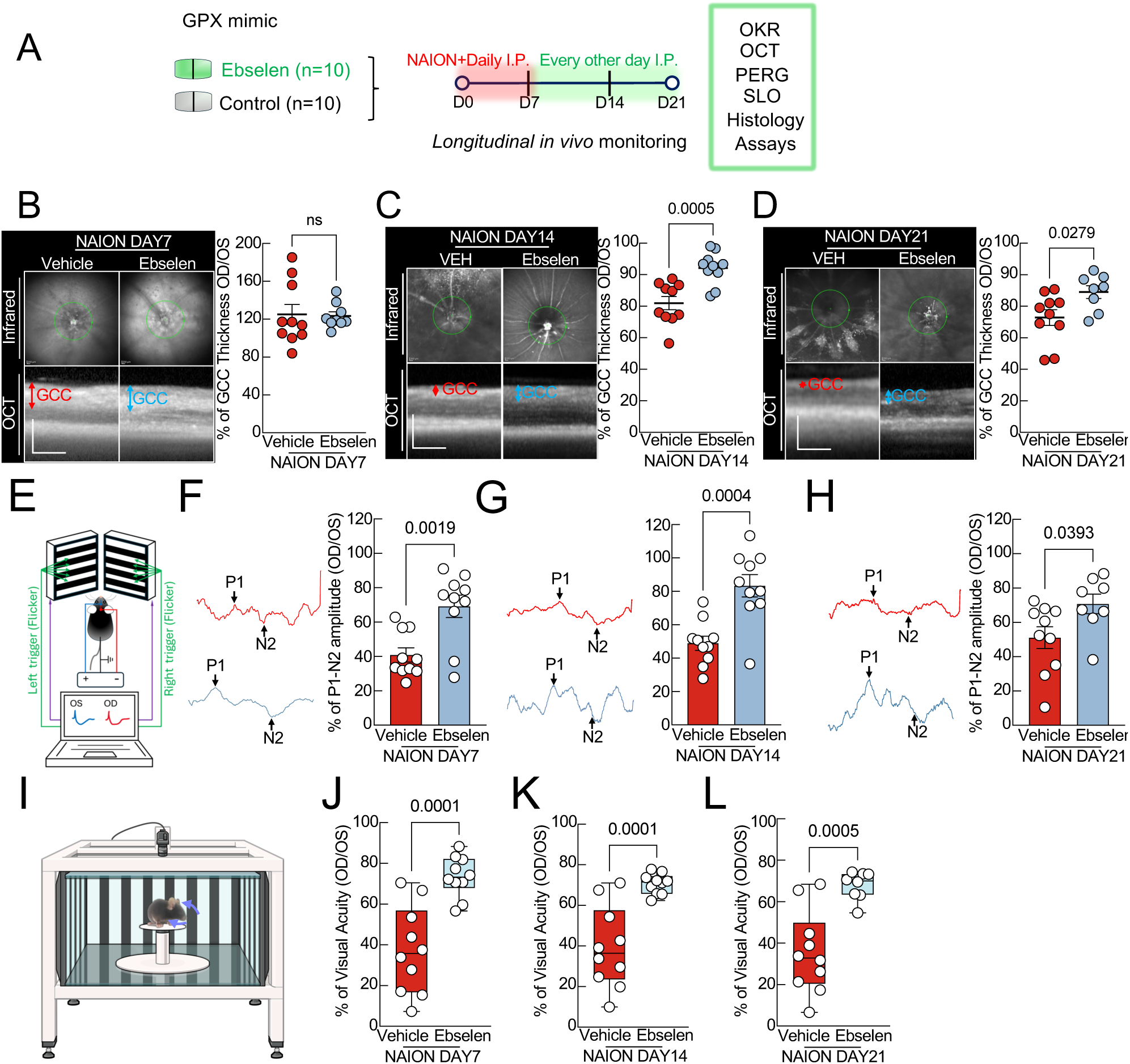
Systemic treatment with GPX-mimic Ebselen significantly preserves visual function in the NAION model. (A) A schematic diagram showing an anti-lipid peroxidation agent, Ebselen was administered intraperitoneally starting on day 1 post-NAION injury, continuing daily through day 7, and subsequently every other day. Ebselen-treated group: n = 10 mice. Vehicle-treated group: n = 10 mice. Longitudinal in vivo monitoring was performed at days 7, 14 and 21 post-NAION injury. (B) Left panels show in vivo OCT images of mouse retinas obtained 7 days after NAION induction. The ganglion cell complex (GCC), comprising the retinal nerve fiber layer to inner plexiform layers, is indicated by the double-headed arrows. Scale bar: 200 *μm*. The right panels present measurements of GCC thickness determined by OCT, with values expressed as percentages relative to the corresponding contralateral eyes. (C) Representative in vivo OCT scans of mouse retinas at 14 days following NAION are displayed on the left. Scale bar: 200 *μm*. The right panels summarize OCT-based measurements of GCC thickness at this time point, normalized to contralateral control eyes and presented as percentages. (D) Left panels depict OCT images acquired from mouse retinas 21 days after NAION injury. The right panels show quantitative analysis of GCC thickness at day 21, expressed as a percentage of the thickness measured in the contralateral eyes. Scale bar: 200 *μm*. One OCT dataset in the Ebselen-treated group at day 7 could not be obtained because a mouse developed transient cataract following anesthesia. Control group: n = 10 mice at days 7, 14, and 21 post-NAION. Ebselen group: n = 9 mice at day 7, n = 10 mice at day 14, and n = 8 mice at day 21. Two mice died before the day 21 analysis. (E) A diagrammatic drawing shows the stimulation of PERG. (F) Left panels display example PERG traces. The right panels show quantification of PERG P1–N2 amplitudes measured 7 days after NAION induction, expressed relative to the corresponding contralateral eyes. Control (n = 10 mice) and Ebselen-treated (n = 10 mice) groups are included. (G) Representative PERG recordings are shown on the left. On the right, P1–N2 amplitudes obtained at day 14 following NAION are summarized and normalized to values from the contralateral control eyes. Control (n = 10 mice) and Ebselen (n = 10 mice). (H) Example PERG waveforms are presented on the left. The right panel summarizes P1–N2 amplitudes recorded 21 days after NAION, reported as a percentage relative to the fellow eyes. Control (n = 10 mice) and Ebselen (n = 8 mice). (I) Visual function was further evaluated by OKR testing to determine visual acuity (Created in BioRender. Yang, M. (2026) https://BioRender.com/yea6tsq). (J) OKR performance at day 7 post-NAION is quantified and presented relative to the contralateral eyes. Control (n = 10 mice) and Ebselen-treated animals (n = 10 mice). (K) Quantitative analysis of OKR at day 14 after NAION, normalized to contralateral control eyes. Control (n = 10 mice) and Ebselen (n = 10 mice). (L) OKR measurements obtained 21 days after NAION induction are summarized and expressed relative to the corresponding control eyes. Control (n = 10 mice) and Ebselen (n = 8 mice). All quantitative results shown in B, C, D, F, G, and H are presented as mean ± s.e.m., and statistical significance was assessed using a two-tailed t-test.

**Fig. 7.**
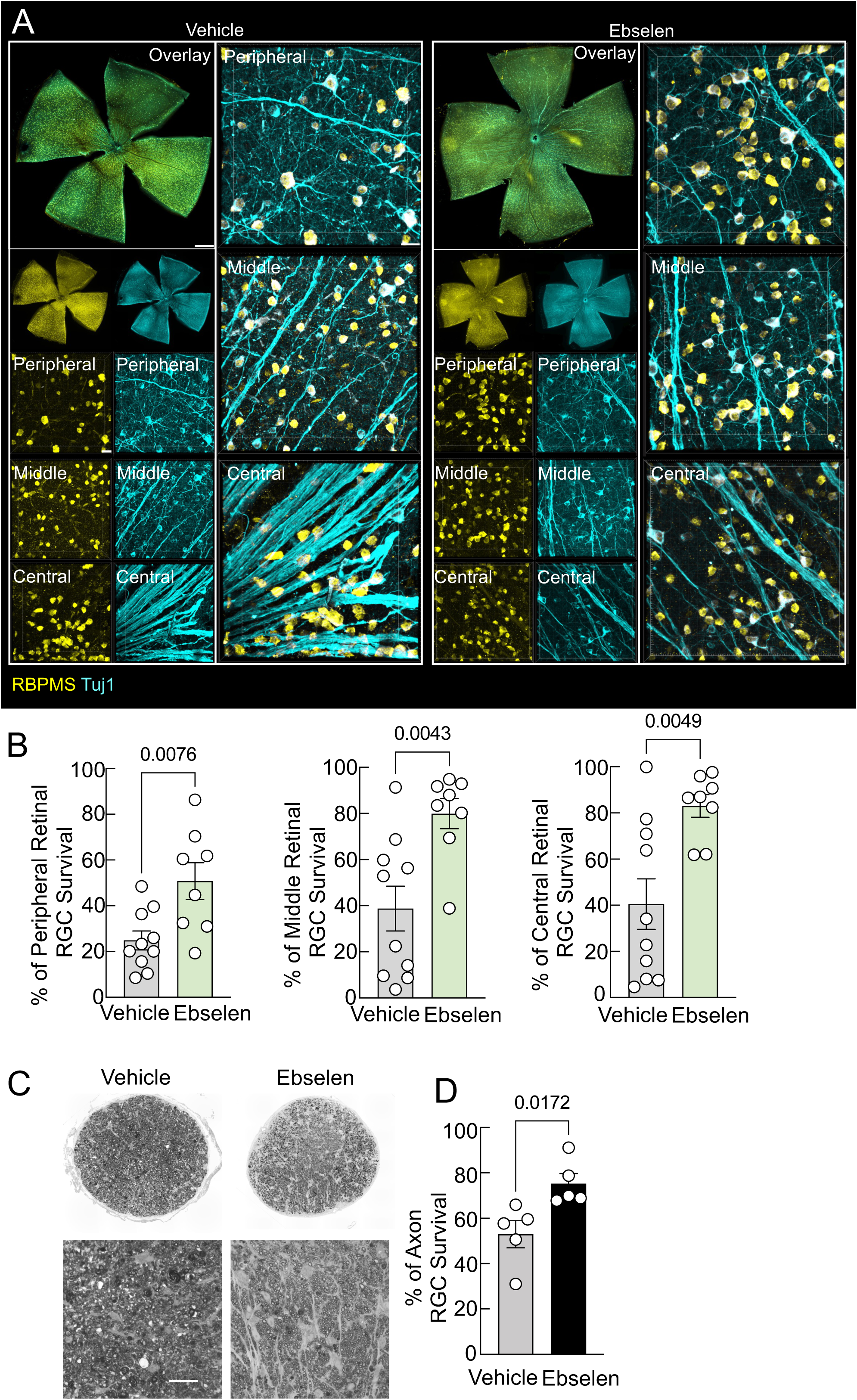
Ebselen promotes significant neuroprotection in both RGC soma and axon in the NAION model. (A) Whole-mount retinal images illustrating RBPMS⁺ and Tuj1⁺ retinal ganglion cells (RGCs) remaining 21 days after NAION induction. Low-magnification views of entire retinas are shown (scale bar, 250 µm), together with higher-magnification images from peripheral, mid-peripheral, and central retinal regions (scale bar, 20 µm). Control group: n = 10 mice; Ebselen-treated group: n = 8 mice. (B) Density of RBPMS⁺ RGCs measured in peripheral, middle, and central retinal areas. Values are expressed as the proportion of surviving RGCs in NAION eyes relative to their corresponding contralateral untreated eyes. (C) Representative light micrographs of paraphenylenediamine-stained semi-thin cross sections of the optic nerve (ON) obtained 21 days after NAION induction (scale bar, 20 µm). (D) Quantitative analysis of axonal preservation in ON semi-thin sections. Axon numbers are normalized to the contralateral control nerves and presented as percentages. Control (n = 5 mice) and Ebselen (n = 5 mice). Data shown in (B) and (D) are reported as mean ± s.e.m., and statistical comparisons were performed using a two-tailed t-test.

We next used intravitreal CTB tracing to assess the integrity of the retinohypothalamic, retinogeniculate, and retinocollicular pathways at 21 days after injury (Fig. S5A). NAION markedly impaired anterograde axonal transport, reducing CTB labeling in the SCN, LGN, and SC to approximately 37% (Fig. S5B), 40% (Fig. S5C), and 62% (Fig. S5D) of uninjured control levels, respectively. Ebselen treatment substantially preserved retinal projections to these central targets, restoring CTB labeling to approximately 76% in the SCN, 71% in the LGN, and 73% in the SC relative to uninjured controls (Fig. S5A–D). These findings indicate that Ebselen preserves long-range visual pathway connectivity by maintaining axonal integrity and anterograde transport from the retina to the brain. Collectively, these results indicate that Ebselen’s GPX-mimetic antioxidant activity reduces oxidative injury, thereby sustaining retinal structure, function, and connectivity after NAION.

### Ebselen treatment reestablishes mitochondrial homeostasis and curbs lipid peroxidation in NAION

To decipher the ROS status of Ebselen-mediated neuroprotection, we labelled mitochondria by intravitreal injection of MitoTracker Green, and the mitochondrial superoxide (O2.^-^) indicator MitoSOX (31) and performed in vivo imaging of these key mitochondrial biomarkers. We found that without treatment, NAION precipitates a profound depletion of mitochondria (Fig. 8A,C) and a concomitant decrease in mitochondrial superoxide flux (Fig. 8B,D) at day 7. Remarkably, treatment with Ebselen dramatically improves axonal and retinal MitoTracker and mitochondrial superoxide label, indicating preservation of axonal and retinal mitochondrial homeostasis. This was accompanied by amelioration of the optic nerve ischemia-activated increase in retinal lipid peroxidation, as indicated by key biomarker MDA (Fig 8E). These findings demonstrate successful Ebselen-mediated inhibition of lipid peroxidation and restoration of mitochondrial homeostasis in a mouse NAION model.

**Fig. 8.**
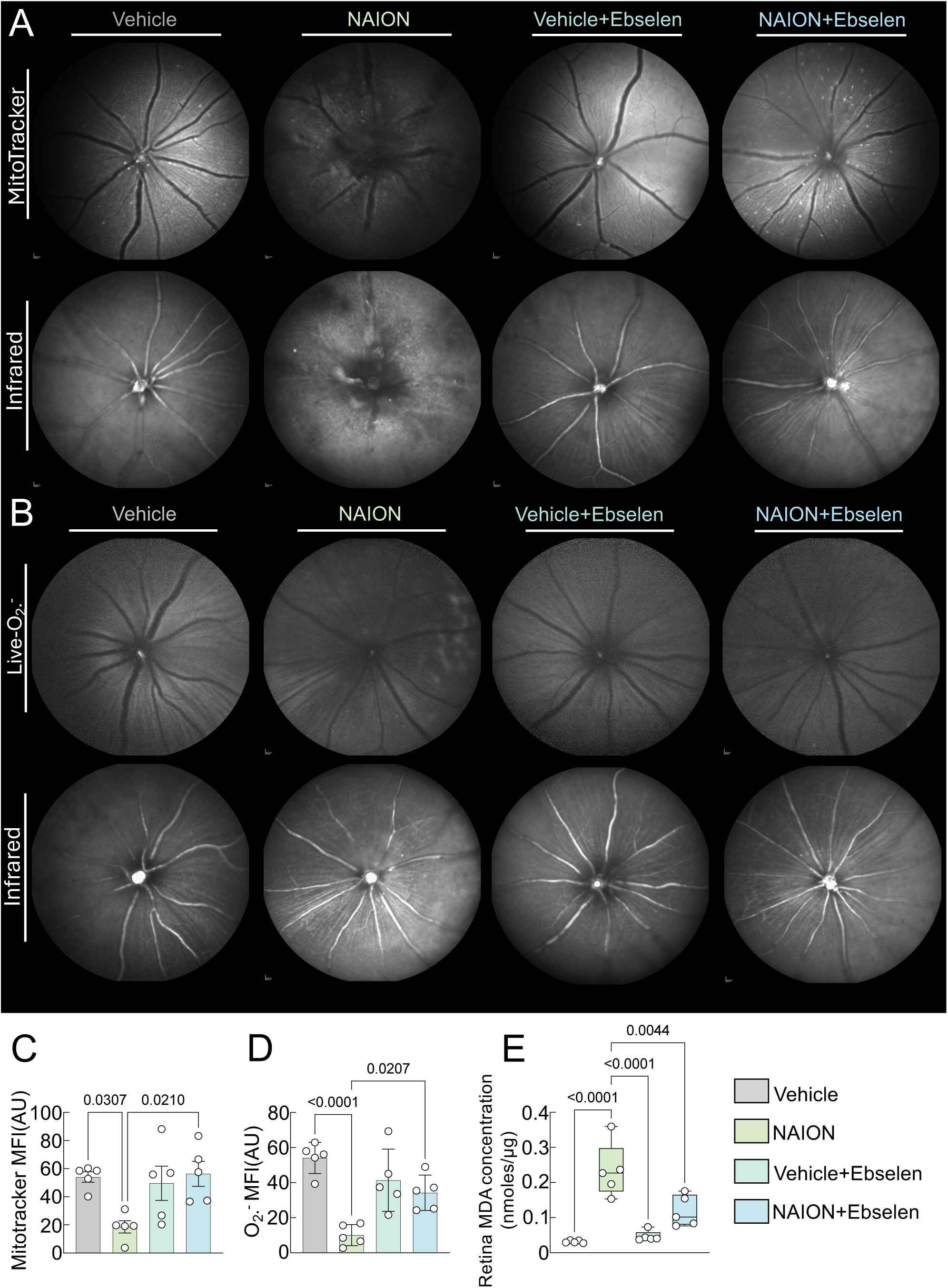
Ebselen improves mitochondrial homeostasis and ameliorates lipid peroxidation in RGC after NAION. (A) Mitochondria abundance in the RGC were monitored by intravitreal injection of Mitotracker in vehicle (n = 5 mice), NAION (n=5 mice), vehicle+Ebselen (n = 5 mice), and NAION+Ebselen (n = 5 mice) groups. (B) Superoxide in the RGC was monitored by intravitreal injection of MitoSOX in Vehicle (n = 5 mice), NAION (n = 5 mice), vehicle+Ebselen (n = 5 mice), and NAION+Ebselen (n = 5 mice) groups. (C) Quantification of mean fluorescent intensity (MFI) of Mitotracker in different groups. (D) Quantification of mean fluorescent intensity (MFI) of superoxide in different groups. (E) MDA levels of the retina in Vehicle, NAION, vehicle+Ebselen, and NAION+Ebselen, were determined. Vehicle (n=5 mice), NAION (n = 5 mice), vehicle+Ebselen (n = 5 mice), and NAION+Ebselen (n = 5 mice). Data are presented as means ± s.e.m, one-way ANOVA with Tukey’s multiple comparisons test.

## DISCUSSION

We show that accumulation of lipid peroxidation products, including malondialdehyde, and 4-hydroxynonenal (4-HNE) in the retinas of NAION patient and animal model. We demonstrate that genetic and pharmacologic augmentation of GPX4 confers robust neuro- and axonal protection and preserves visual function in experimental NAION. Crucially, RGC- and mitochondria-specific GPX4 overexpression markedly preserves RGC survival, axonal integrity, and central visual projections. Treatment of animal model of NAION using Ebselen, a GPX-mimetic that has been tested in clinical trials of stroke, hearing loss, and tinnitus, recapitulates these RGC protective effects by restoring mitochondrial redox balance and normalizing reactive oxygen species dynamics. Leveraging real-time multiparametric in vivo imaging to directly interrogate axonal and retinal metabolism and function, we show that without treatment, NAION induces a profound collapse in RGC axonal energy state as shown by in vivo ATP imaging, accompanied by dysregulated reactive oxygen species signalling, as seen on in vivo mitochondrial imaging, and dramatic alteration of the retinofugal projections to the brain, including to the lateral geniculate nucleus, the major projection to the primary visual cortex conferring vision. Mechanistically, we postulate that GPX4-mediated detoxification of lipid peroxides preserves axonal and retinal mitochondrial abundance and function, thereby sustaining the energetic demands required for axonal transport and neuronal survival in NAION. Translationally, these findings define a plausible pathway toward clinically meaningful interventions for NAION using GPX4 as a novel target (fig. S6).

We observed robust activation of lipid peroxidation in human and experimental NAION, as evidenced by accumulation of 4-hydroxynonenal, malondialdehyde, and oxidative DNA damage. These findings place membrane lipid oxidation downstream of ischemic injury but upstream of irreversible neuronal degeneration, positioning lipid peroxidation as a critical mechanistic link between vascular insult and axonal failure. There is strong indirect evidence that the key NAION risk factors such as cardiovascular disease(40), atherosclerosis(41), and aging(42, 43) operate in a milieu of increased lipid peroxidation, even though direct lipid peroxidation measurements in NAION patients are not yet well characterized. Although ferroptosis is classically defined as an iron-dependent process(44), we did not detect significant iron accumulation in this NAION model, noting that any iron dysregulation may be transient and temporally restricted; this finding suggests that lipid peroxidation–driven neurodegeneration in NAION is driven primarily by ischemia-induced oxidative stress and mitochondrial dysfunction rather than sustained iron overload, thereby broadening the conceptual framework of ferroptosis-related injury in the optic nerve.

A defining feature of NAION is the selective susceptibility of the anterior optic nerve (45), a region with exceptionally high mitochondrial density, metabolic demand, and lipid-rich axonal membranes (23). These characteristics, combined with structural constraints of the optic nerve head, render RGC axons uniquely vulnerable to ischemia-induced oxidative stress (18). Occlusion of the blood supply to the anterior optic nerve has never been demonstrated in NAION (46). This may be because NAION is not simply a disease of vascular insufficiency but rather by a failure to terminate lipid peroxidation–driven membrane damage at the level of the axon and mitochondria. Also, lipid peroxidation is a mechanistically distinct form of oxidative injury. It cannot be mitigated by conventional reactive oxygen species scavenging(47). Once initiated, phospholipid hydroperoxides propagate membrane damage and bioenergetic collapse through self-amplifying reactions. This necessitates dedicated enzymatic detoxification rather than generic antioxidant approaches. We propose NAION is driven by pathological lipid oxidation, and therapeutic success requires direct detoxification of phospholipid hydroperoxides.

Accumulating experimental and clinical evidence implicates oxidative stress as a convergent pathological driver across diverse RGC-degenerative conditions, including glaucoma (48), hereditary optic atrophy (49), and ischemic and traumatic optic neuropathies (50), but it is not yet studied in NAION context. In these settings, lipid peroxidation not only acts downstream of primary insults such as ischemia, inflammation, or elevated intraocular pressure, but also amplifies neurodegeneration through feed-forward interactions with mitochondrial dysfunction, impaired axonal transport, and defective cellular stress responses. Accordingly, elucidating the interplay between oxidative stress, retinal ganglion cell bioenergetics, and neuronal survival is critical for the rational design of neuroprotective strategies in optic nerve disease, and our findings provide proof-of-concept that targeting lipid peroxidation represents a viable therapeutic approach for NAION.

As a novel therapeutic target for NAION, GPX4 occupies a singular position in cellular redox defence as the only enzyme capable of directly detoxifying phospholipid hydroperoxides within biological membranes (25, 51, 52). Loss of GPX4 activity is sufficient to trigger ferroptotic cell death (23), whereas its preservation is essential for neuronal survival under oxidative stress. Our data demonstrate that RGC-specific overexpression of GPX4 markedly preserves ganglion cell survival, axonal integrity, and visual function following NAION. Beyond structural preservation, GPX4 overexpression restored fundamental bioenergetic and redox homeostasis within RGC axons. and implicated in membrane destabilization, protein adduct formation, cytoskeletal disruption, lysosomal dysfunction, and inflammatory signalling in age-related macular degeneration(1), glaucoma(26), cataract(53), and retinal aging (54–56). Our study in NAION is relevant to other optic neuropathies such as glaucoma(26). RGCs possess exceptionally high metabolic demands and a high risk of lipid peroxidation due to their long axons and prolonged unmyelinated segments within the retina, rendering them critically dependent on mitochondrial oxidative phosphorylation for sustained ATP production (14, 57). In this study, we provide proof-of-concept that GPX4-based gene therapy can be neuroprotective for NAION. Although gene therapy is classically considered a treatment for monogenic disorders, gene therapy targeting GPX4 can be a viable therapy for the treatment of NAION, given the accessibility of the ocular compartment for gene therapy and relatively reduced off-target effects.

Notably, mitochondrial-targeted GPX4 provided slightly greater protection than the cytosolic isoform, highlighting mitochondria as a key site where lipid peroxidation and metabolic failure converge during ischemic optic nerve injury. In line with these observations, MitoGPX4 exhibited greater neuroprotective efficacy than CytoGPX4 in the NAION model, suggesting that mitochondrial targeting may represent a more effective neuroprotective strategy. Importantly, protection extended beyond the retina to central visual pathways. MitoGPX4 overexpression preserved anterograde transport to the lateral geniculate nucleus and superior colliculus, indicating that axonal continuity and functional connectivity within the visual system are maintained. This systems-level preservation is particularly relevant for NAION, where axonal degeneration and transport failure are primary determinants of permanent vision loss. Notably, impaired axonal transport is also a central pathogenic feature in glaucoma (58, 59) and optic neuritis(60, 61), where disruption of long-range connectivity contributes directly to visual dysfunction. Thus, preservation of mitochondrial integrity and axonal transport may represent a broadly applicable neuroprotective strategy for maintaining visual pathway function across optic neuropathies.

Complementing genetic intervention, we demonstrate that Ebselen, a clinical trial tested(34), lipophilic glutathione peroxidase mimetic, recapitulates many of the protective effects of GPX4 overexpression. Ebselen preserved RGC structure and function, mitigated lipid peroxidation, and restored mitochondrial integrity and ROS balance after NAION. These results provide proof-of-concept that pharmacologic reinforcement of the GPX4-glutathione axis can deliver neuroprotection without gene therapy, strengthening the translational relevance of our findings. In contrast to gene therapy, small-molecule pharmacological approaches like Ebselen are readily applicable to treatment of age-related diseases like NAION, which can be optimized to allow for flexible dosing and timing for clinical translations. Ebselen exhibits remarkable redox-modulatory and glutathione peroxidase–like activities in ameliorating oxidative stress. The current limitations of Ebselen is its hydrophobicity and challenges of systemic administration, which may necessitate higher doses and increase the risk of off-target effects. Further reformulation and drug development are needed to improve ocular delivery strategies, including enhanced solubilization to maximize retinal and optic nerve bioavailability.

Collectively, our study suggests NAION as a disorder of ischemia-induced lipid peroxidation-driven neurodegeneration and establishes GPX4 as an essential determinant of RGC resilience to ischemic injury. By demonstrating efficacy across human tissue, cell-type-specific genetic rescue, mitochondrial functional restoration, and pharmacologic intervention, we provide a strong translational foundation for ferroptosis-targeted therapies. Future work will focus on optimizing delivery strategies, dosing, durability, and safety, with the ultimate goal of advancing GPX4-based or GPX4-mimetic therapies toward clinical evaluation for NAION and related optic neuropathies.

## MATERIALS AND METHODS

### Animals and Husbandry

C57BL/6J wild-type mice of both sexes were sourced from Jackson Laboratory. Upon arrival, animals were housed at Stanford University School of Medicine under regulated conditions: a constant 12-hour light/dark schedule, ambient temperature of 25 ± 2°C, and relative humidity maintained between 40% and 60%. Animals had unrestricted access to food and water throughout the study. All procedures involving animal subjects were vetted and approved by the Stanford University IACUC in accordance with institutional regulations.

### Plasmid Construction and Recombinant AAV Generation

We have previously published generation of AAV2 vectors for RGC-specific, mitochondria and cytoplasmic-targeted GPX4 overexpression (26). These isoforms share an identical C-terminal sequence that encompasses the catalytic active site, yet diverge at the N-terminus, which harbors subcellular targeting motifs that dictate distinct intracellular localizations (62). As a selenoprotein, GPX4 requires the SECIS element in the 3′ untranslated region (UTR) to translate an in-frame UGA codon as selenocysteine. We targeted the mitochondrial and cytosolic GPX4 forms and engineered AAV vectors to express them under the RGC-specific mSncg promoter (63), incorporating SECIS elements to ensure precise selenocysteine incorporation (Fig. S3A). Following intravitreal AAV2 delivery, HA-tagged GPX4 colocalized with RBPMS-positive RGCs, indicating robust RGC-specific expression (Fig. S3B).

To generate the constructs for MitoGPX4 and CytoGPX4, coding sequences (CDS) and their corresponding selenocysteine insertion sequences (SECIS) were amplified from mouse tissue cDNA libraries as described previously (26). PCR amplification was performed using 2×Phanta Flash Master Mix (Vazyme, P520-03) with the specific primers listed in Table 1. The resulting fragments were cloned into the pAM-AAV-mSncg(0.27kb)-3HA-WPRE vector backbone using MfeI, XbaI, MluI, or HindIII restriction sites. The pAAV-hSyanpsin-(cyto)-iATPSnFR2-S29W-A95K-HaloTag plasmid was obtained from Addgene (#209663). All plasmids were isolated and purified using the FastPure EndoFree Plasmid Midi Kit (Vazyme, DC205-01) per the manufacturer’s protocol. AAV particles were produced in HEK293T cells via triple transfection of the target construct, AAV2 wild type and pHelper (Stratagene) using Polyethylenimine reagent (Polysciences, Inc 24765). Cells were harvested 72 hours post-transfection, and viruses were purified via iodixanol gradient centrifugation (64). Viral titers were determined by SDS-PAGE (65). For experimental delivery, 2 μL of the viral suspension was administered via intravitreal injection.

**Table 1.** PCR Primers designed in this study.

### Ophthalmological Procedures and Functional Assessments

Experimental procedures, including intravitreal injections, histological evaluation of the optic nerve (ON) and retinal ganglion cells (RGCs), and various in vivo imaging and functional assays, were performed as previously described (26, 63, 66–71). Briefly, retinal morphology was assessed via optical coherence tomography (OCT) and confocal scanning laser ophthalmoscopy (cSLO). Visual function was quantified using pattern electroretinography(PERG) and optokinetic response (OKR) measurements. Summary protocols for these techniques are detailed below.

### Intravitreal Administration and Viral Transduction

Mice (approximately 6 weeks old) were anesthetized with a ketamine/xylazine cocktail (0.08 mg/g and 0.01 mg/g, respectively) administered according to body weight. To ensure optimal volume delivery, 2 μL of vitreous humor was first aspirated using a polished glass microcapillary pipette inserted at the peripheral retina near the ora serrata. Subsequently, 2 μL of the AAV suspension was injected into the vitreous chamber. A two-week incubation period was maintained post-injection to allow for robust transgene expression. For anterograde axonal labeling, mice received a secondary intravitreal injection of 2 μL CTB-Alexa Fluor 555/647 (2 mg/mL; Invitrogen).

### Pharmacological Intervention

For experiments involving antioxidant treatment, Ebselen (MedChemExpress, HY-13750) was administered to wild-type mice via intraperitoneal (i.p.) injection at a dose of 10 mg/kg, consistent with previously established protocols (72–74). Ebselen was initially dissolved in DMSO to prepare a stock solution, which was aliquoted and stored at −20 °C. On the day of administration, a single aliquot was thawed and diluted in 20% cyclodextrin (MedChemExpress, Cat. No.: HY-101103) to the desired concentration immediately prior to intraperitoneal (IP) injection. The treatment regimen was initiated on the day of NAION induction (Day 0). Mice received daily injections for the first 7 days, followed by a maintenance schedule of one injection every other day for the remainder of the study.

### Non-arteritic anterior ischemic optic neuropathy (NAION) surgery and treatment

The NAION procedure was performed two weeks post-intravitreal injection of adeno-associated virus (AAV) or before the Ebselen injection, when the mice were aged 6-7 weeks. To induce NAION (75–79) (fig. S1A), mice are first anesthetized with an intraperitoneal injection of ketamine/xylazine, and their pupils are dilated with 1% tropicamide. A glass coverslip with artificial tears (Systane Ultra Lubricant Eye Drops, Alcon Laboratories, Fort Worth, TX) is applied to the cornea to permit visualization of the fundus. Induction of ischemia is achieved through photochemical thrombosis: a photosensitizing dye, Rose Bengal (1.29 mM), is administered via retro-orbital injection to the OS eye, followed immediately by targeted laser illumination of the optic nerve head (ONH) to the OD eye. A 532 nm green laser is used to deliver approximately 50 mW of power over a 400 μm diameter spot for a series of 15 pulses. This interaction generates localized superoxide radicals, causing capillary thrombosis and subsequent optic nerve edema. Post-procedural success is confirmed by the presence of optic disc swelling and pallor, typically monitored via OCT, OKR, and PERG (fig.S2B-D). After the procedure, the cornea was coated with a neomycin-containing ophthalmic ointment (Akorn, Somerset, NJ) to protect the ocular surface and promote postoperative recovery.

### Measurement of malondialdehyde (MDA) level

Retinal malondialdehyde (MDA) concentrations were quantified using a colorimetric Lipid Peroxidation Assay Kit (ab118970, Abcam) following the manufacturer’s protocol. Briefly, two freshly harvested mouse retinas per sample were homogenized on ice in MDA lysis buffer containing butylated hydroxytoluene (BHT). Insoluble debris was removed via centrifugation at 13,000×g for 10 minutes at 4°C. Supernatants were collected, and standard curves were generated using MDA concentrations ranging from 0 to 5 nmol/well. After equilibrating reagents to room temperature, samples and standards were reacted with a developer solution at 95 °C for 60 minutes to form MDA-TBA adducts. Following a 10-minute cooling period on ice, 200 µL of each reaction mixture was transferred to a 96-well plate, and absorbance was recorded at 532 nm. To ensure accurate comparison, MDA levels were normalized to total protein content as determined by a bicinchoninic acid (BCA) protein assay.

### Quantification of Retinal Iron Homeostasis

Total iron, ferrous iron (Fe²⁺), and ferric iron (Fe³⁺) levels were determined using an Iron Assay Kit (ab83366, Abcam) following the manufacturer’s instructions. Freshly harvested retinas were homogenized on ice in the provided iron assay buffer and clarified via centrifugation (13,000 × g) for 10 min at 4 °C in a refrigerated microcentrifuge to remove insoluble material.

Supernatants were collected and processed alongside a standard curve (0–10 nmol/well). To measure total iron, an iron reducer was added to convert all Fe^3+^to Fe^2+^, while samples for ferrous iron measurement were processed without the reducer. Following a 30-minute incubation at 37°C, 100 µL of the Iron Probe was added to each well, and the plate was incubated in the dark for 60 minutes at 37°C. Absorbance was recorded at 593 nm. Ferric iron levels were calculated by subtracting the ferrous iron concentration from the total iron concentration. All values were normalized to total protein content as measured by a BCA assay.

### *In vivo* morphometry via spectral-domain optical coherence tomography (SD-OCT)

In vivo retinal architecture was evaluated using spectral-domain optical coherence tomography (SD-OCT) with a 30° lens (Heidelberg Engineering) as previously described (70, 80). High-resolution images were acquired in central scan mode, with 100 frames averaged per scan to optimize the signal-to-noise ratio. The thickness of the ganglion cell complex (GCC), comprising the retinal nerve fiber layer (RNFL), ganglion cell layer (GCL), and inner plexiform layer (IPL), was quantified using commercially available segmentation software (Heidelberg engineering, Germany) as described previously(66). To assess neurodegeneration, the average GCC thickness in the peripapillary region of the injured eye was normalized to the contralateral control eye and expressed as a percentage. All imaging and subsequent analyses were performed by researchers masked to the experimental groups.

### *In vivo* Confocal Scanning Laser Ophthalmoscopy (cSLO) imaging

For in vivo cSLO imaging in mice, the procedure begins with deep anesthesia and full pupillary dilation using 1% tropicamide and 2.5% phenylephrine. To compensate for the high refractive power of the rodent eye and prevent corneal desiccation, a custom 10D contact lens is applied to the cornea. The mouse is positioned on a 3D-adjustable platform to align the pupillary axis with the cSLO camera, which utilizes 488 nm laser wavelengths to perform point-by-point raster scans of the retina. By adjusting the confocal aperture and internal focus, the retinal ganglion cell layer fluorescence images are captured and averaged in real-time to minimize noise and motion artifacts. For MitoSOX (Invitrogen, M36006) imaging, 10 μl DMSO was used to dissolve the MitoSOX powder, and further diluted in 1:10 ratio for intravitreal injection 1 hour before imaging. For MitoTracker (Invitrogen, M7514) imaging, 75 μl DMSO was used to dissolve one tube of MitoTracker power, and further diluted in 1:10 ratio for intravitreal injection 1 hour before imaging. For in vivo ATP imaging, AAV-iATPSnFR2 was intravitreally injected to the mice 14 days before NAION and imaging. For quantification of mean fluorescence intensity, each in vivo fluorescence images of the retina were captured for each mice across all experimental groups. The immunofluorescence intensity of the whole filed of view was measured using Fiji/ImageJ software, following background subtraction.

### Electrophysiological Assessment of Retinal Function

Pattern electroretinography (PERG) was performed using the Miami PERG system (Intelligent Hearing Systems, Miami, FL) following previously described protocols (26, 70, 80). To ensure physiological stability, mice were placed on a feedback-controlled heating pad maintained at 37°C. Corneal clarity was preserved throughout the procedure using Systane lubricant drops. For signal acquisition, a three-electrode configuration was employed: a reference electrode was placed subcutaneously at the interaural midline, a ground electrode at the tail base, and a common active needle electrode at the snout for simultaneous binocular recording. Visual stimuli were delivered via two 14 × 14 cm LED panels positioned 10 cm from each eye. The stimulus consisted of a black-and-gray reversing pattern (spatial frequency: 0.052 cycles/degree; 85% contrast; 800 cd/m^2^ luminance). Signals were recorded using asynchronous binocular acquisition, with a final waveform generated by averaging two consecutive blocks of 100 and 300 traces. RGC function was quantified by the P1–N2 amplitude (the voltage difference between the first positive peak at ∼100 ms and the subsequent negative trough). The P1–N2 amplitude of the experimental eye was normalized to the contralateral control eye, and all data were analyzed by investigators blinded to the treatment groups.

### Behavioral Assessment of Visual Acuity

Spatial visual acuity was quantified via the optokinetic response (OKR) using the OptoMotry system (Cerebral Mechanics Inc.), as previously described (26, 67, 70). Unrestrained mice were positioned on a central pedestal within a virtual reality arena composed of four 17-inch LCD monitors. An overhead camera monitored the animal’s behavior. A virtual cylinder displaying vertical sine wave gratings was rotated at a fixed speed; clockwise rotation specifically stimulated the left eye, while counterclockwise rotation stimulated the right eye. Testing began at a baseline spatial frequency of 0.1 cycles/degree. If a mouse exhibited distinct head-tracking movements in the direction of the rotation during a 5-second stimulus interval, the spatial frequency was incrementally increased. The threshold for spatial acuity was defined as the highest frequency at which the mouse reliably demonstrated tracking behavior. To ensure data integrity, all assessments were performed during the morning to account for circadian variability. Results are presented as the percentage of acuity in the experimental eye relative to the contralateral control. All measurements were conducted by masked investigators.

### Tissue Clearing and Anterograde Projection Mapping

To visualize the central projections of RGC axons, we utilized an adapted iDISCO (immuno-labeled localized three-dimensional intact-tissue-clearing) protocol (81). Forty-eight hours following intravitreal CTB-555 administration, mice were transcardially perfused with 4% paraformaldehyde (PFA). The brain and attached optic nerves (ON) were carefully dissected and stabilized in a 1% agarose block. The agarose-embedded samples underwent a multi-day clearing sequence: serial dehydration in methanol (20%, 40%, 60%, 80%, and 100% in PBS; 1 day each), followed by delipidation in dichloromethane (DCM)/methanol (2:1; 1 day), and final refractive index matching in 100% DCM (1 day) and dibenzyl ether (DBE; 1 day). Cleared samples were mounted on a spike holder and imaged using a Light-Sheet Ultramicroscope II. The tissue was illuminated bi-directionally by six 3.89 µm light sheets (NA 0.149). Images were captured through a 2x objective with a 0.63× zoom using 561 nm and 640 nm diode lasers. Z-stacks were acquired at 6 µm intervals. Finally, optical sections were processed into maximum intensity projections using Fiji/ImageJ to visualize the complete axonal trajectory.

### Histological Processing and Quantitative Morphometry

To assess retinal health and neurodegeneration, we utilized standardized immunohistochemical techniques for both whole-mount and cross-sectional preparations. The protocols for retinal tissue processing and immunostaining were adapted from previously established methods (66, 82, 83). Following transcardiac perfusion with 4% PFA and overnight cryoprotection in 30% sucrose, retinal whole-mounts were isolated and processed for triple-labeling. Tissues were blocked for 30 minutes in 10% normal goat serum with 2% Triton X-100 and subsequently incubated overnight at 4°C with primary antibodies against RBPMS (1:4000; ProSci) to identify RGC somata, Tuj1 (1:500; Biolegend, 801202) for axonal integrity, and rat anti-3HA (1:500; Novus, NBP2-50416) to confirm construct expression.

Secondary detection was performed using species-specific Alexa Fluor antibodies (568, 488, or 647; 1:200 dilution; Jackson ImmunoResearch) for 1 hour at room temperature. Following rigorous washing protocols (3 × 30 minutes in PBS), retinas were mounted with Fluoromount-G and imaged using a Zeiss LSM 880 confocal microscope and a Keyence BZ-X800 system. RGC survival was determined by sampling 6–9 randomized fields (332×332 µm) spanning the central, middle, and peripheral retinal regions. RBPMS-positive cells were quantified via Fiji/ImageJ by masked investigators, with survival rates expressed as a percentage relative to the contralateral control eye.

For a detailed analysis of the retinal architecture and oxidative stress markers, mouse eyeballs were embedded in Tissue-Tek OCT, snap-frozen on dry ice, and sectioned using a Leica cryostat. These cross-sections were immunostained with anti-RBPMS (1:4000; ProSci) to identify RGC somata, anti-Tuj1 (1:500; Biolegend, 801202) for axonal integrity, anti-8-OXOG (1:500; QED Bioscience, 12501)and anti-4-Hydroxynonenal (1:500; Alpha Diagnostics, HNE51-5) to evaluate lipid peroxidation levels. Secondary antibodies, including Alexa Fluor 594- or 647-conjugated goat anti-rabbit, 488-conjugated goat anti-guinea pig, and 594-conjugated goat anti-mouse, were applied at a 1:200 dilution for 1 hour. Quantitative assessment of protein expression and lipid peroxidation was achieved by capturing three immunofluorescence images per sample across triplicate independent experiments using the Keyence BZ-X800 microscope. The mean fluorescence intensity (MFI) within the ganglion cell layer (GCL) was meticulously calculated using Fiji/ImageJ software. All measurements incorporated background subtraction to ensure objective comparisons of oxidative stress and structural preservation across the various experimental cohorts.

### Optic Nerve Histology and Axon Quantification

To evaluate axonal preservation, optic nerve semi-thin sections were prepared and analyzed according to established protocols. Transverse sections of 1 µm thickness were cut using a Leica EM UC7 ultramicrotome at a distance of 2 mm distal to the globe (1.5 mm from the injury site) and stained with 1% para-phenylenediamine (PPD) in a 1:1 methanol mixture to provide high-contrast visualization of the myelin sheaths. The entire cross-section of each nerve was imaged and stitched using a Keyence microscope under a 100× objective to provide a complete high-resolution topographical map. For quantification, a systematic random sampling approach was implemented using the “AxonCounter” plugin in ImageJ (68), where a 10 µm × 10 µm grid was applied to approximately 10% of the total nerve area. Within these frames, axons were manually enumerated using the ImageJ cell counter tool, and the resulting mean counts were normalized to the contralateral uninjured control to determine the percentage of surviving axons. To ensure the objectivity of the results, all histological assessments were conducted by investigators blinded to the experimental groups.

### Statistical Analyses

All statistical analyses and data visualization were performed using GraphPad Prism (version 10) or R software. Data are expressed as the mean ± standard error of the mean (SEM). For comparisons between two independent groups, a two-tailed unpaired Student’s t-test was employed. When analyzing differences among multiple groups, we utilized one-way or two-way analysis of variance (ANOVA), followed by appropriate post hoc multiple comparison tests to determine specific group differences. The threshold for statistical significance was pre-defined as p<0.05.

## Abbreviations

4-HNE: 4-hydroxynonenal
cSLO: confocal scanning laser ophthalmoscopy
GCC: ganglion cell complex
GPX4: Glutathione peroxidase 4
MDA: malondialdehyde
NAION: nonarthritic anterior ischemic optic neuropathy
OCT: optical coherence tomography
OKR: optokinetic response
PERG: pattern electroretinogram
PUFAs: polyunsaturated fatty acids
RGCs: retinal ganglion cells

## Data Availability Statement

The data supporting the findings of this study are available within the article and its supplemental information.

## Acknowledgment

We thank the Liao lab members for critical discussion. M.Y. is supported by 2024 ARVO Foundation Early Career Clinician-Scientist Research Award. We thank Stanford University Cell Sciences Imaging Facility (CSIF (RRID:SCR_017787)) for the Light-Sheet Ultramicroscope serice. We thank Prof.Yang Hu from Stanford Ophthalmology for the generous support of the mSncg AAV Vectors. The authors acknowledge the use of large language model (LLM) editing tools, including ChatGPT 5.3 and Gemini Pro, for writing assistance. All scientific content, interpretations, and conclusions were generated and verified by the authors.

## Author Contributions

M.Y., Y.P.L.J designed the experiments. M.Y., S.M, J.P. and C.J.P established the methodology and M.Y., S.M, J.P. and C.J.P were involved in sample collection and quantification. X.B.R. produced AAVs. M.Y., S.M,., J.P., A.A., A.S., T.A. H.W., and Y.P.L.J prepared the manuscript with support from all the authors.

## Supplementary Materials

**fig. S1.**
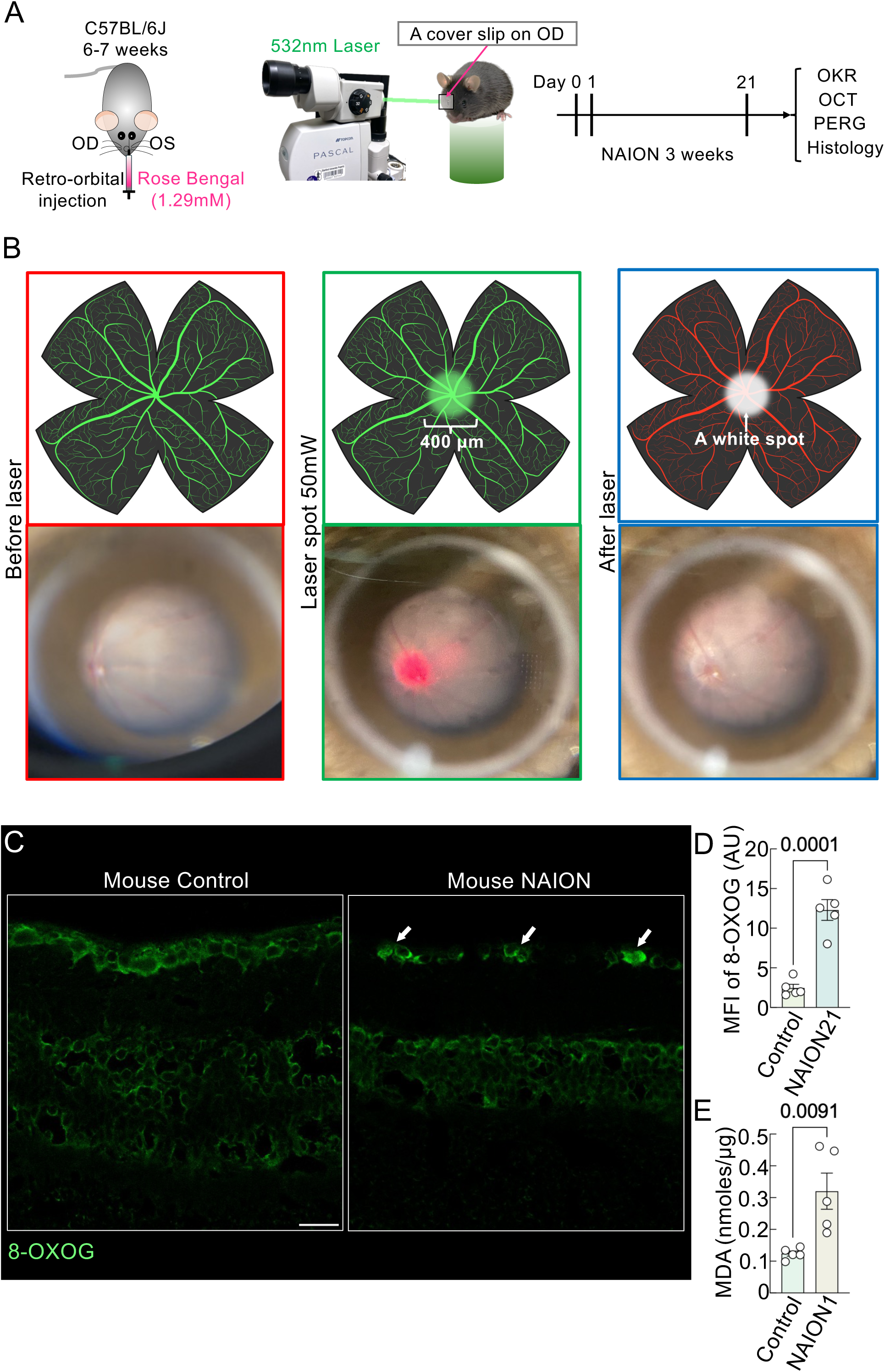
Increased 8-Oxoguanine (8-OXOG) in surviving retinal ganglion cells in mouse nonarthritic anterior ischemic optic neuropathy (NAION). (A) Workflow of the experimental NAION model. To induce NAION, mice were anesthetized with intraperitoneal ketamine/xylazine and pupils dilated using 1% tropicamide. A coverslip with artificial tears was placed on the cornea to enable fundus visualization. Ischemia was induced by photochemical thrombosis: Rose Bengal (1.29 mM) was delivered via retro-orbital injection, followed by targeted illumination of the optic nerve head with a 532 nm laser (∼50 mW, 400 μm spot, 15 pulses). (B) The fundus changes of the mouse NAION model. A white spot appears in the center of the mouse after ischemia was induced by photochemical thrombosis. Figure created with BioRender.com and modified by the authors (Created in BioRender. Yang, M. (2026) https://BioRender.com/2gfn1v2). (C) Representative confocal micrographs of mouse retinal sections stained for 8-OXOG. Scale bar, 20 µm. (D) Mean fluorescence intensity of 8-OXOG was measured in vehicle-treated and NAION mice 21 days after laser-induced injury. Data are shown as mean ± s.e.m.; **p < 0.01, two-tailed t-test. N=5 control or NAION eyes. (E) Retinal malondialdehyde (MDA) levels were measured in vehicle and NAION mice at day 1 post-NAION injury. Each point represents one sample derived from two pooled retinas. Vehicle (n = 5 mice, 10 retinas) and NAION (n = 5 mice, 10 retinas).

**fig. S2.**
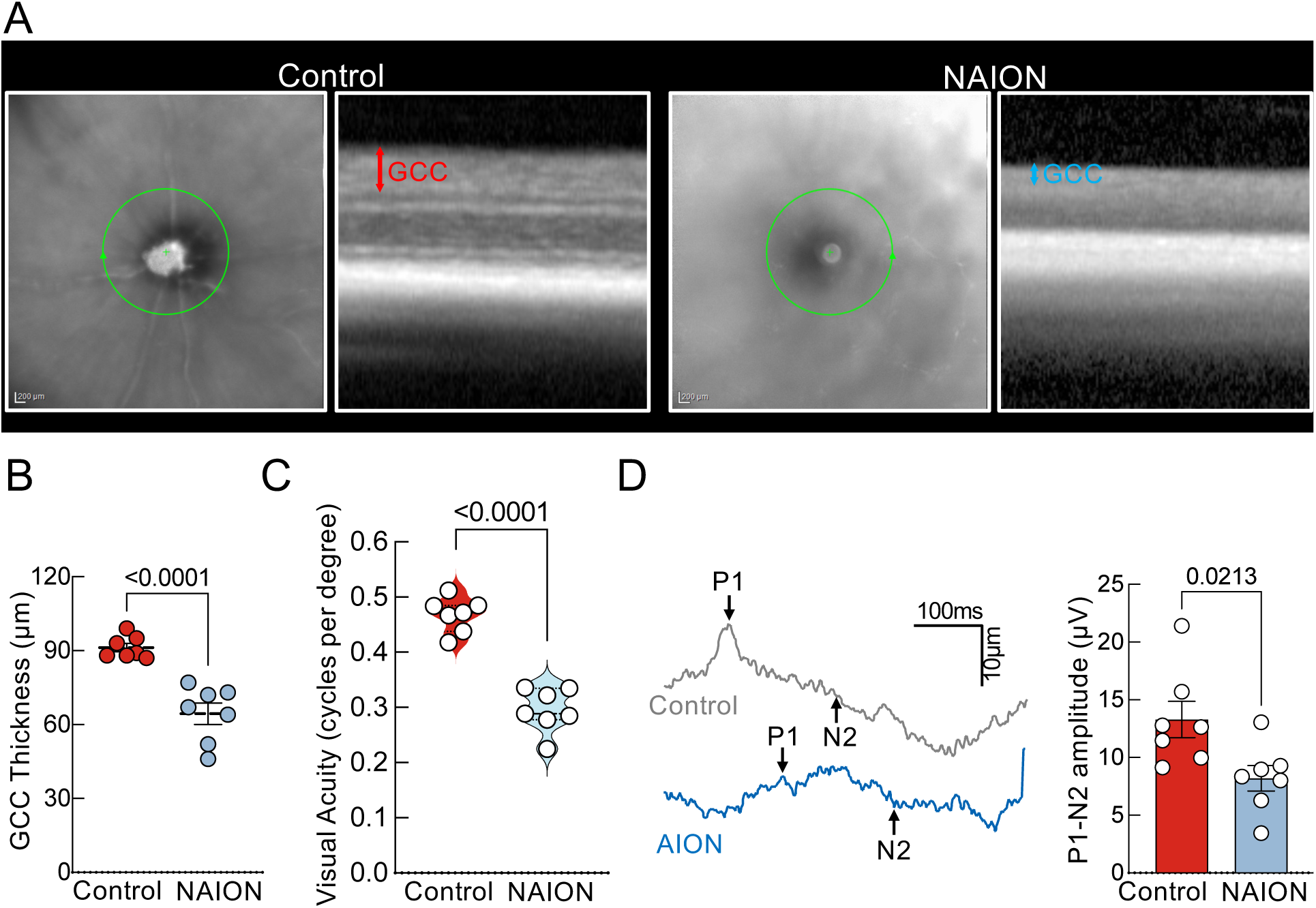
Robust structural and functional deficits confirm successful induction of mouse NAION. (A) Representative in vivo OCT images of mouse retinas at day 21 after NAION injury. GCC: ganglion cell complex, including retinal nerve fiber layer to inner plexiform layers; indicated by double-end arrows. (B) GCC thickness was evaluated using OCT 21 days after NAION induction. Measurements from NAION eyes were normalized to those from the corresponding contralateral eyes and expressed as percentages. Control (n = 7 mice) and NAION (n = 7 mice). (C) Visual function was assessed by the OKR at 21 days following NAION. Results are presented relative to the contralateral control eyes and expressed as percentages. Control (n = 7 mice) and NAION (n = 7 mice). (D) Left panels show representative PERG traces. The right panels summarize the P1–N2 amplitudes of PERG recorded 21 days after NAION induction, normalized to contralateral eyes and displayed as percentages. Control (n = 7 mice) and NAION (n = 7 mice).

**fig. S3.**
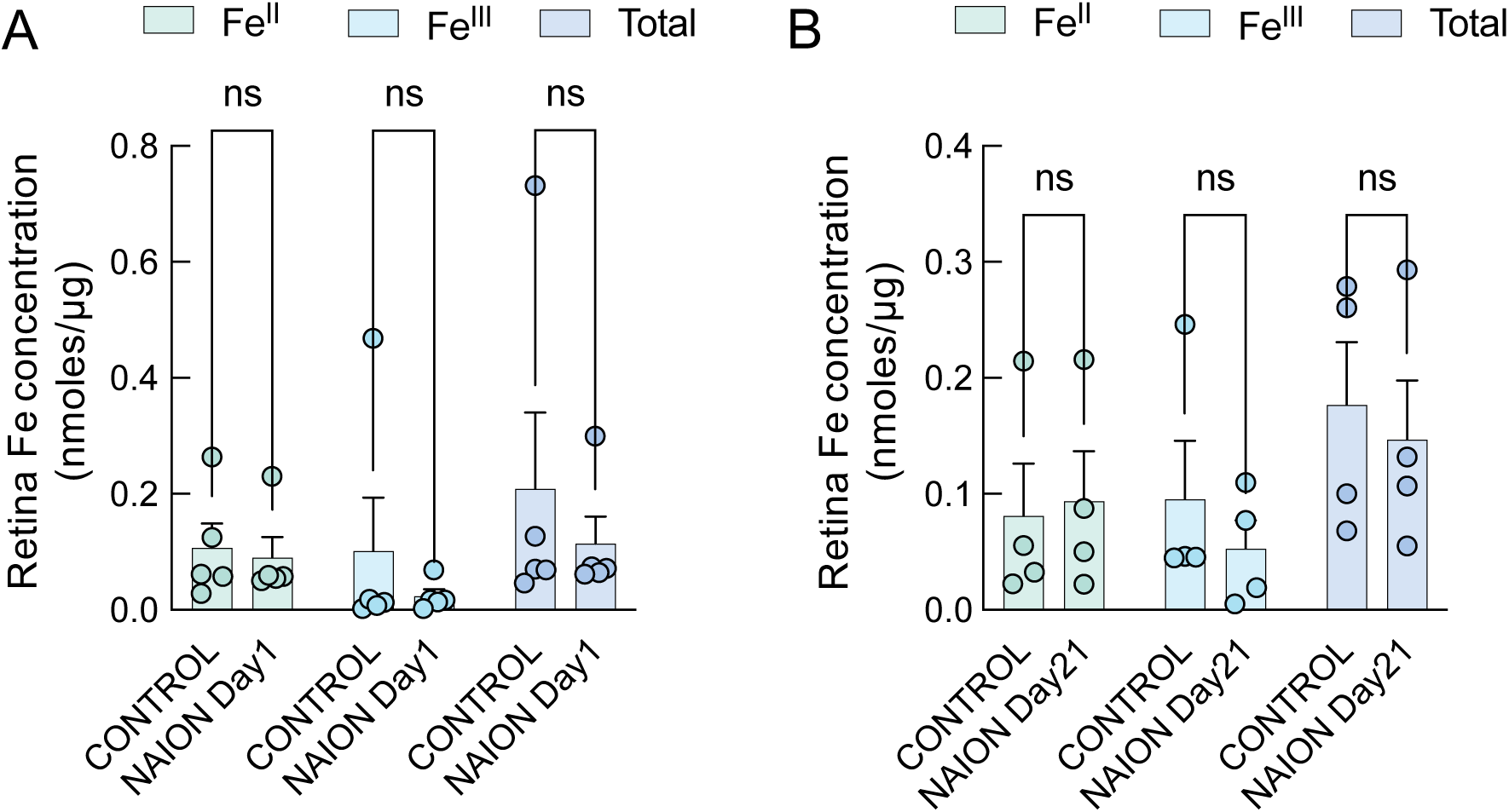
Iron level is not significantly altered in an experimental model of NAION. (A) Total iron, ferric iron, and ferrous iron in control and day 1 post-NAION injury. (B) Total iron, ferric iron, and ferrous iron in control and day 21 post-NAION injury.

**fig. S4.**
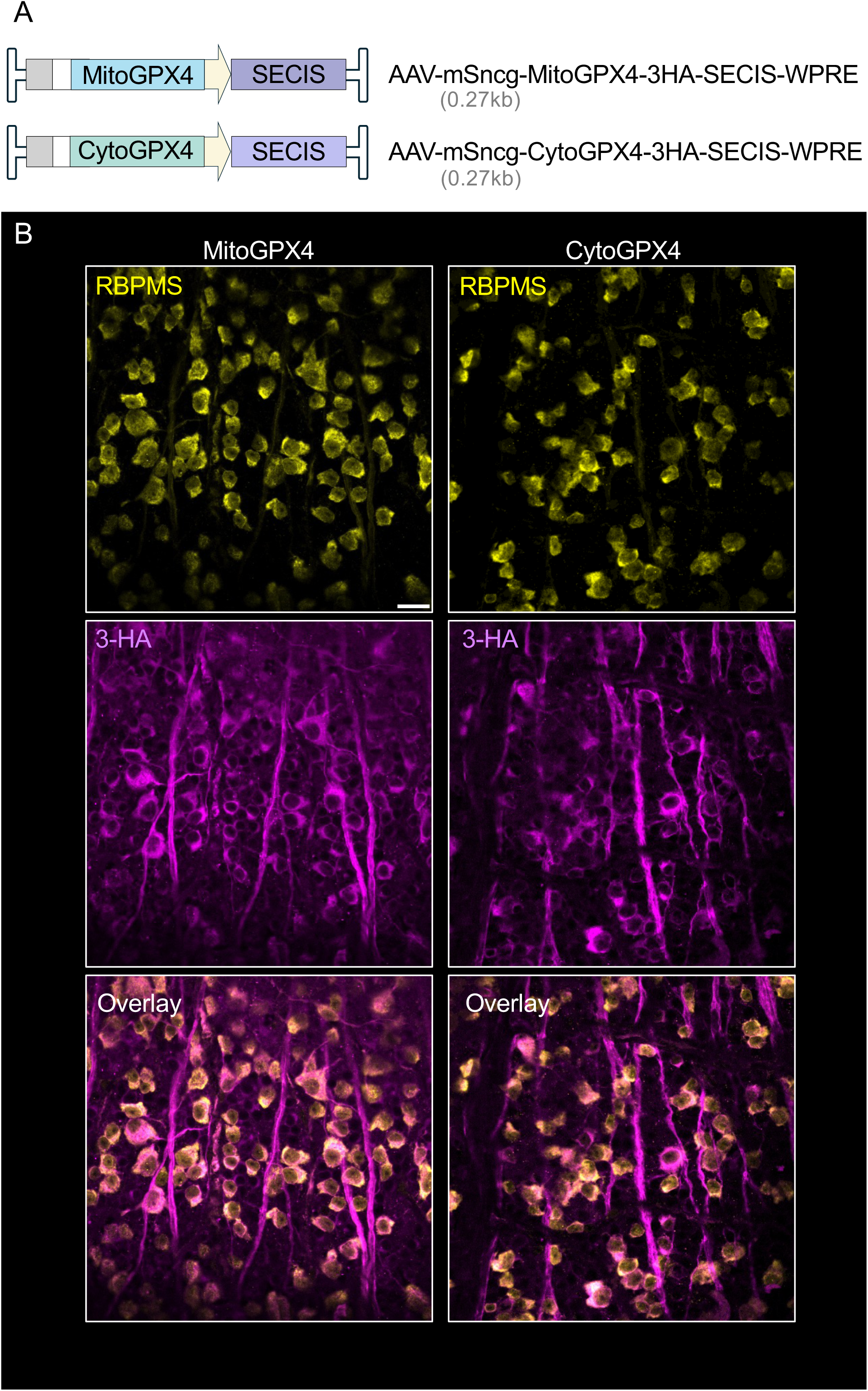
RGC-specific overexpression of MitoGPX4 and CytoGPX4. (A) Schematic of AAV2 vectors expressing mitochondrial GPX4 (MitoGPX4) or cytosolic GPX4 (CytoGPX4) carrying a 3×HA tag and SECIS (selenocysteine insertion sequence) under the control of the mSncg promoter. (B) AAV-mediated RGC-specific transgene expression detected by HA immunostaining 2 weeks after injection intravitreally. Scale bar, 20 µm.

**fig. S5.**
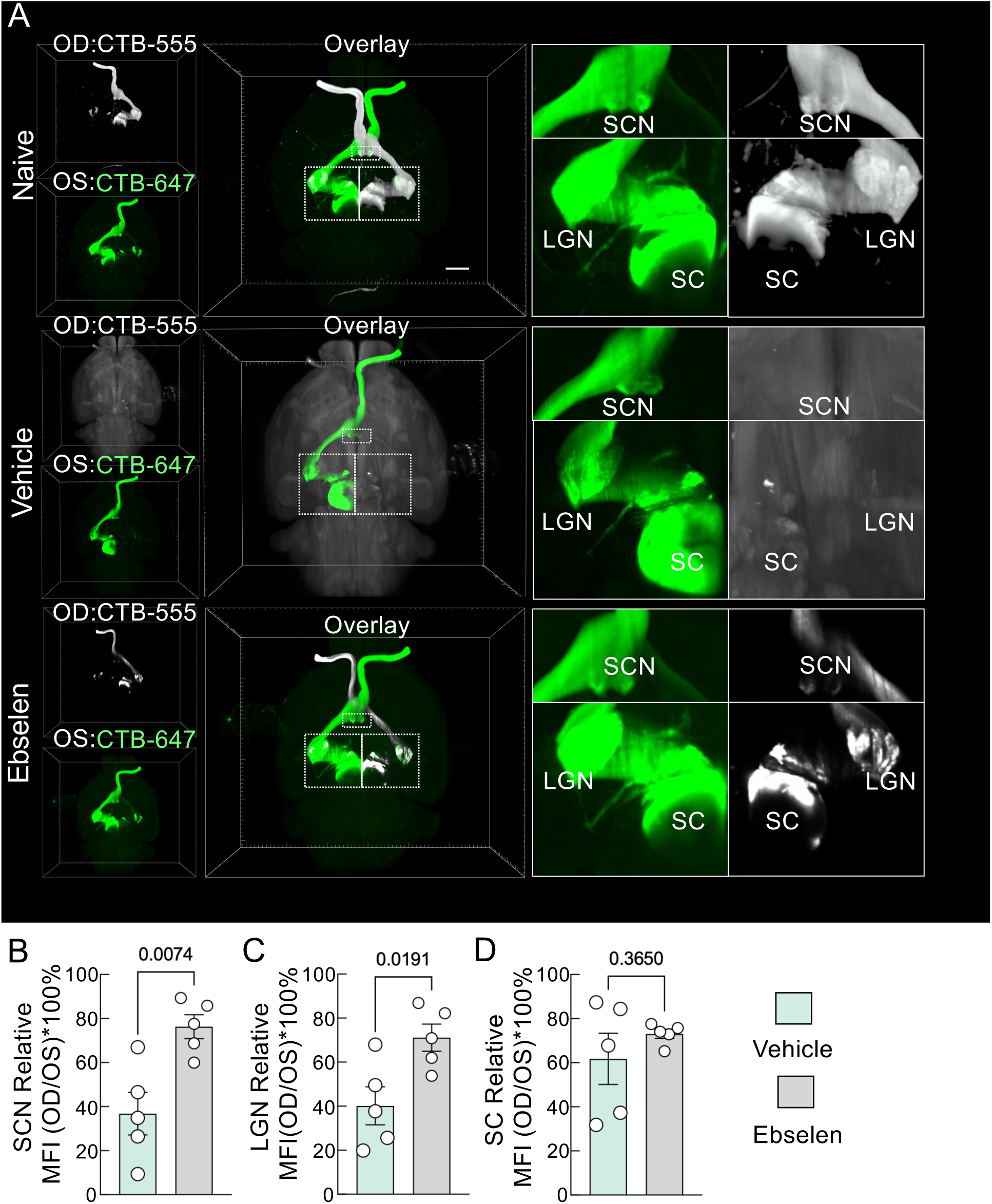
Ebselen provides substantial neuroprotection across the visual pathway in a NAION model. (A) Representative 3D images of the eye and brain were reconstructed following cleared-tissue imaging using a Light-sheet Ultramicroscope II. The visual transportation is labeled by intravitreal injection of cholera toxin B (CTB)-555 and CTB-647 to each eye. (B) Quantification of relative fluorescent intensity of CTB signal in superior colliculus nucleus (SCN), Vehicle (n = 5 mice), Ebselen (n = 5 mice). (C) Quantification of relative fluorescent intensity of CTB signal in the lateral geniculate nucleus (LGN), Vehicle (n = 5 mice), Ebselen (n = 5 mice). (D) Quantification of relative fluorescent intensity of CTB signal in the SC, Vehicle (n = 5 mice), Ebselen (n = 5 mice). Scale bar: 1000 µm. Data are presented as means ± s.e.m, two-tailed t-test.

**fig. S6.**
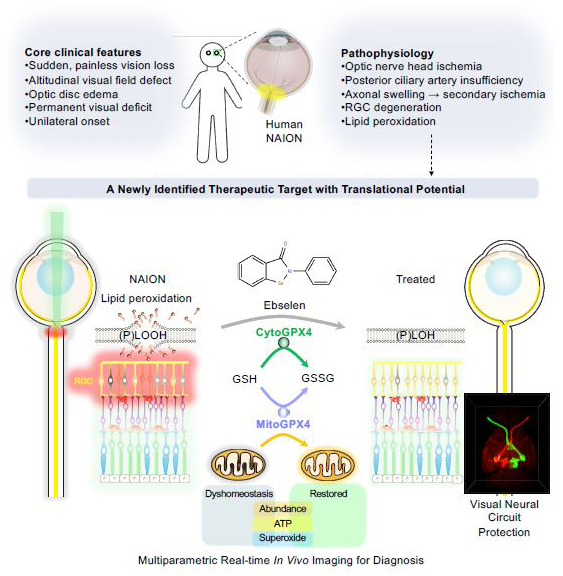
Diagram depicting the mechanisms by which GPX4 and Ebselen inhibit lipid peroxidation. This study identifies lipid peroxidation as an essential mechanism linking ischemic injury to retinal ganglion cell degeneration in NAION. By integrating human patient tissue with a rigorously validated mouse model, we show that cell-type-specific augmentation of GPX4, the only enzyme that directly detoxifies membrane phospholipid hydroperoxides, preserves neuronal survival, axonal integrity, mitochondrial bioenergetics, and visual function. Importantly, these protective effects are recapitulated by the clinically tested GPX-mimetic Ebselen, establishing lipid peroxidation-targeted GPX4 modulation as a disease-modifying and translationally actionable therapeutic strategy for NAION. (Created in BioRender. Yang, M. (2026) https://BioRender.com/xlsc9nz)

**Movie 1. The induction of the mouse NAION model.**

**Movie 2. 532nm laser-exposed mouse retina with retro-orbital injection of Rose Bengal.**

**Movie 3. 532nm laser-exposed mouse retina without retro-orbital injection of Rose Bengal.**

**Movie 4. Mouse optokinetic responses (OKR) in NAION + Capsid, NAION + CytoGPX4, and NAION+MitoGPX4 groups.**

**Movie 5. Representative 3D video of the naïve visual pathway was reconstructed following cleared-tissue imaging using a Light-sheet Ultramicroscope II.**

**Movie 6. Representative 3D video of the NAION visual pathway was reconstructed following cleared-tissue imaging using a Light-sheet Ultramicroscope II.**

**Movie 7. Representative 3D video of the NAION+MitoGPX4 visual pathway was reconstructed following cleared-tissue imaging using a Light-sheet Ultramicroscope II.**

**Movie 8. Mouse optokinetic responses (OKR) in vehicle + NAION and Ebselen + NAION groups.**

